# Harvesting effects on forest condition indicators across Iberian forests: Implications for the EU Nature Restoration Regulation

**DOI:** 10.64898/2025.12.11.693644

**Authors:** Pedro Rebollo, Paloma Ruiz-Benito, Enrique Andivia, Miguel A. Zavala, Julen Astigarraga, Susanne Suvanto, Verónica Cruz-Alonso

**Affiliations:** Departamento de Biodiversidad, Ecología y Evolución, Facultad de Ciencias Biológicas. Universidad Complutense de Madrid, Madrid, España; Universidad de Alcalá. Grupo de Ecología y Restauración Forestal, Departamento Ciencias de la Vida. Facultad de Ciencias, Alcalá de Henares; Universidad de Alcalá, Grupo de Investigación en Teledetección Espacial, Departamento de Geología, Geografía y Medio Ambiente, Alcalá de Henares; Department of Physical Geography and Ecosystem Science, Lund University, Lund, Sweden; Natural Resources Institute (Luke). Helsinki, Finland

**Keywords:** Carbon stocks, Forest Inventory, Harvesting intensity, Native species, Standing deadwood, Structural diversity, Tree species diversity

## Abstract

1. Forest degradation is posing a growing threat to biodiversity conservation, climate regulation and nature contributions to people worldwide. The EU Nature Restoration Regulation (NRR) recognise the need of healthy forests and has established indicators to assess the condition of forest ecosystems. Forest management practices, especially harvesting may contribute to improve these indicators by, for example, reducing tree density and promoting tree species diversification.
2. Here, we assessed temporal trends in forest condition indicators (hereafter, indicators) across Iberian forests since the 80s and evaluated how harvesting occurrence and intensity modulated these trends. Using 46,354 plots from the Spanish Forest Inventory (1986–2022), we analysed trends in indicators depending on stand diversity (monospecific or mixed), protection status (protected or unprotected), origin of the stand (natural or planted), and biogeographical region (Mediterranean or temperate).
3. Overall, indicators increased over time. Harvesting occurrence reduced the increases in aboveground carbon stocks, structural diversity, tree species diversity, and standing deadwood; however, it contributed to increase the proportion of native species in specific forest types. Medium to high harvesting intensity negatively impacted aboveground carbon stocks, while medium intensities increased tree species diversity but reduced the structural diversity.
4. *Synthesis and applications.* Our results suggest that indicators are increasing as stand develops in absence of disturbances such as harvesting. Tree harvesting cannot be considered as a silver bullet to achieve the objectives of the NRR, but it can contribute under certain conditions – specifically at low intensities for carbon stocks and at medium intensities for species diversity. The naturally positive trend of indicators underlines the need to establish thresholds values and minimum rates of changes that distinguish restoration outcomes from natural dynamics. Finally, our study also highlights the key role of forest inventories in monitoring forest condition over time and across diverse landscapes.

## 1. INTRODUCTION

The Nature Restoration Regulation (NRR) aims to restore 20% of degraded ecosystems in Europe by 2030 and all ecosystems in need of restoration by 2050. As a part of this effort, European countries are required to increase biodiversity and enhance ecosystem functioning (European Union, 2024). The NRR places biodiversity recovery at the forefront of policy and land management across member states, while facing climate change and adaptation to global change risks. Forests, which cover approximately 43% of the EU extension, play a fundamental role in sustaining that biodiversity and consequently nature contributions to people such as climate regulation and carbon sequestration (Díaz et al., 2018; European Union, 2024). Assessing the condition of forest ecosystems is therefore essential to plan restoration efforts and achieve the objectives of the NRR.

The NRR defines different forest condition indicators, including carbon stocks, the proportion of forests with uneven-aged structures, tree species diversity, share of forests dominated by native species and the amount of standing deadwood, among others (European Union, 2024). These indicators are used to assess forest degradation or deviation from the NRR’s objectives and to evaluate the effectiveness of restoration measures. Carbon sequestration is a key forest function, playing a pivotal role in climate change mitigation by regulating the global carbon cycle and storing over 45% of terrestrial carbon (FAO, 2025). Enhancing forest carbon stocks directly supports the European Union’s climate mitigation objectives, as outlined in the EU Climate Law and the *“Fit for 55”* package to reduce carbon emissions by 55% by 2030 (European Comission, 2021). The NRR explicitly aligns restoration targets with these climate goals, reinforcing the role of forest ecosystems in achieving net-zero emissions. Tree species diversity and structural complexity can enhance the resilience of forest ecosystems to climate change and disturbances, while together with deadwood can provide habitat for multiple species (Gamfeldt et al., 2013; Heidrich et al., 2020; Rebollo et al., 2025). By contrast, forest plantations associated to monospecific stands of non-native species and managed as short rotation crops, led to low biodiversity and soil degradation (Castro-Díez et al., 2019; Wohlgemuth et al., 2022), although they can provide higher levels of carbon sequestration (Lázaro-Lobo et al., 2025). Thus, forest condition indicators proposed by the NRR not only assess ecosystem functioning, but also monitor conservation outcomes and inform adaptative management measures (Burrascano et al., 2021; Corona et al., 2011). However, the effectiveness of measuring these indicators at large spatial extents, such as those covered by the NRR depends on the availability of consistent, spatially explicit and long-term datasets that allow for reliable monitoring of forests dynamics (Moreno-Fernández, Oliveira, et al., 2025; Paillet et al., 2024).

National forest inventories provide a robust framework for assessing forest condition, thanks to their extensive spatial coverage and standardised data collection protocols. These attributes allow for the detection of temporal trends in forest functioning across varied ecological contexts (Tomppo et al., 2010; Vidal et al., 2016). Forest inventories have inherent limitations such as relatively short temporal coverage, the likelihood of underrepresentation of extreme events because data is not continuously monitored over space and time, and the focus on the vegetation layer (Moreno-Fernández, Breidenbach, et al., 2025; Ruiz-Benito et al., 2020; Serra-Maluquer et al., 2025). Despite these limitations, forest inventories have been widely used for studying forest ecosystem dynamics and represent a critical tool to assess the effect of climate change on forest ecosystems (Sagarin & Pauchard, 2010, 2012). In fact, the NRR explicitly recommends using forest inventories to quantify forest condition indicators (European Union, 2024). Although forest inventories have great potential, they are still rarely used for identifying the drivers of forest functioning in restored landscapes (but see Cruz-Alonso et al., 2019). Integrating the availability of high-quality forest inventory data with policy actions could strengthen the scientific foundation of restoration policies, such as the NRR and other landscape-scale conservation initiatives.

Human use of forest resources shapes ecosystem structure and dynamics, conditioning the capacity of forests to act as carbon sinks, reservoirs of biodiversity and to provide ecosystem services (Daigneault et al., 2022; Duncker et al., 2012; Triviño et al., 2023). Specifically, tree harvesting interventions might reduce canopy density and promote regeneration, which can ultimately enhance forest diversity, structural complexity and the provision of ecosystem services (D’Amato et al., 2011; Li et al., 2020). Consequently, forest functioning and condition largely depends on the frequency and intensity of harvesting (Nabuurs et al., 2013; Paillet et al., 2010). Forest inventory data are helping to characterise these regimes across Europe (Suvanto et al., 2025), yet we still know little about how harvesting frequency and intensity affect forest functioning over time and at large spatial extents. Thus, quantifying the impact of prevailing harvesting practices on forest condition indicators is key, as in some cases it could enhance certain indicators (i.e. carbon stocks; Zhang et al., 2024).

The Mediterranean basin is a biodiversity and climate change hotspot (Lazoglou et al., 2024; Myers et al., 2000), shaped by a long historical human-forest relationship (Blondel, 2006). Mediterranean forests are highly vulnerable to degradation due to historical high land-use pressures and increasing water stress, leading to low productivity compared to temperate ones (Penca & Tănăsescu, 2025). The human footprint has created complex landscapes with forests displaying varying degrees of naturalness. In Spain, the afforestation and reforestation policies led to the increase in planted forest areas during the twentieth century mainly composed by fast-growing species (Vadell et al., 2016). The subsequent and frequent lack of management over decades has produced dense and homogeneous planted stands (Ruiz-Benito et al., 2012), which can lead to increased tree decline and more intense disturbances (Rebollo et al., 2024; Repeto-Deudero et al., 2025). Monospecific natural or planted stands are preferred for timber production due to operational simplicity, but mixed-species forests have shown greater resilience and productivity under stress conditions compared to monospecific stands (Pretzsch & Schütze, 2021; Ureña-Lara et al., 2025). Beyond the inherent complexity of forest landscapes, conservation objectives have become increasingly central to forest management in recent decades. For example, the Spanish Law 42/2007 of Natural Heritage and Biodiversity restricts harvesting in protected areas due to conservation and disturbance prevention purposes. The variety of climates, forest management regimes and the vulnerability of Mediterranean forests, makes Spain a particularly valuable region for studying the consequences of tree harvesting on forest condition, as well as for testing the potential of forest inventories to support monitoring of restoration success.

Here, we investigated forest condition at large spatial extent using the Spanish Forest Inventory (hereafter SFI) to calculate NRR indicators and quantified how harvesting influences the temporal trends of the indicators across different Iberian forests. Specifically, we examined: (i) the temporal trends in the indicators; (ii) how these trends differed based on harvesting occurrence; and (iii) how harvesting intensity affected indicator trends across forest types defined by stand diversity (monospecific or mixed), protection status (protected or unprotected), origin of the stand (planted or natural), and biogeographical region (Mediterranean or temperate). To characterise the indicators, we used 46,354 SFI plots with two or three consecutive surveys between 1986 and 2022. We hypothesised that the effects of tree harvesting on the indicators would vary depending on the indicator studied and the forest type. We expected aboveground carbon stocks and structural diversity to decrease with harvesting intensity, and tree species diversity to peak at medium harvesting intensity, consistent with the intermediate disturbance hypothesis (Connell, 1978). Evaluating the consequences of tree harvesting on forests and its alignment with restoration goals is critical to ensure the effective implementation of the NRR.

## 2. METHODOLOGY

### 2.1. Study area and National Forest Inventory data

The study area covers the large edaphoclimatic gradient of the Iberian Peninsula, from oceanic climates in the north to semi-arid climates in the southeastern region (De Castro et al., 2005), and contrasting soil types composed of siliceous, limestone and clay soils (Instituto Geográfico Nacional, 2019). Different forest types are widely distributed throughout the study area (Fig. S1.1), being *Quercus ilex* L., *Quercus suber* L., and *Quercus pyrenaica* Wild., the most abundant angiosperm species and *Pinus halepensis* Mill., *Pinus pinaster* Ait., and *Pinus sylvestris* L. as the main gymnosperm species (MITECO, 2023).

To assess the temporal trends of the indicators in Iberian forests, we used data from the Spanish Forest Inventory. This dataset encompasses a network of plots systematically distributed across areas with more than 5% forest cover, arranged on a 1-km^2^ cell grid (Villaescusa & Díaz, 1998). Each SFI plot consists of four circular subplots arranged concentrically in a nested design depending on the diameter at breast height of the sampled trees (hereafter d.b.h.): 5 m radius for trees with 7.5-12.4 cm of d.b.h.; 10 m radius for 12.5-22.4 cm of d.b.h.; 15 m radius for 22.5-42.4 cm of d.b.h. and 25 m radius for > 42.4 cm of d.b.h. For each sampled tree, tree d.b.h and height, species identity and status (alive, dead present, dead absent) are recorded among other variables. Three comparable censuses of SFI are currently available: 2SFI (1986–1996), 3SFI (1997–2007) and 4SFI (2008–present). In addition, shrub species identity and cover (%) are sampled in the 10 m radius subplots. Although the information of the 4SFI is not available for the entire country, for this study we used: (i) permanent plots with data available for all three censuses (i.e. 2SFI, 3SFI and 4SFI; 20,345 plots) to study the trends over time (objective 1); and (ii) permanent plots between 23SFI, and between 34SFI for the autonomous communities where two surveys were available (objectives 2 and 3), totalling 46,354 plots.

### 2.2. Forest condition indicators calculated with the Spanish Forest Inventory data

We calculated five indicators from the seven proposed in the NRR using SFI data: carbon stocks, structural diversity, tree species diversity, dominance of native tree species and standing deadwood. As the NRR refers to carbon stocks in terms of organic carbon in litter and mineral soil (European Union, 2024), we first explored the overall temporal trends of the litter and soil related variables available in the SFI (see variable description in Appendix S2 and exploratory analyses in Fig. S2.1). However, this information is qualitative and insufficient to calculate soil organic carbon stocks. Instead, we estimated belowground and aboveground carbon stocks (Mg C ha^-1^) of living adult trees. Although this approach does not directly measure soil carbon, it provides complementary information on carbon sequestration in biomass, which is also a key indicator of forest condition and directly contributes to the restoration objectives outlined in the NRR (European Union, 2024). Thus, we estimated carbon stocks by calculating tree-level belowground and aboveground biomass from tree d.b.h and height, applying species-specific allometric equations for the main species of the Iberian Peninsula according to Ruiz-Peinado et al. (2011, 2012; Table S1.1). Biomass values were then converted to carbon stocks using species-specific carbon content factors (Montero et al., 2005). When species-specific equations were not available, we used those of closest phylogenetically related species. We restricted carbon stocks analyses to plots where allometric equations were available for all the recorded species.

Structural diversity was characterised as the coefficient of variation of tree size (ratio of the standard deviation to the mean d.b.h.; dimensionless). Tree species diversity was characterised using the Shannon index weighted by basal area, which accounts for both species richness and evenness (dimensionless; Shannon, 1948). Basal area was calculated as the sum of the cross-sectional area of the trunk at breast height for each living sampled individual (m² ha⁻¹). Shrub diversity was similarly assessed using the Shannon index weighted by shrub cover, calculated as the percentage of ground area covered by each shrub species (dimensionless). We calculated the dominance of native tree species as the percentage of basal area of native tree species in each plot respect to total stand basal area (%). We identified the non-native species based on the Spanish catalogue of non-native species (MITECO, 2013; Table S1.2). Finally, standing deadwood (m^2^ ha^-1^) was calculated as the basal area of standing dead trees in the final census that were alive in the initial census (i.e. 2SFI in the 23SFI period and 3SFI in the 34SFI period).

To study the indicator trends, we calculated the data at the plot level (i.e. sum of basal area) and calculated the difference between the final and initial value of each indicator for the studied SFI period (i.e. 23SFI and 34SFI period). As standing deadwood per plot was calculated by comparing two censuses, we calculated the difference between 23SFI information and 34SFI information to study the temporal trend. In addition, detailed data on shrub species was only available for the 4SFI census, therefore, temporal trends could not be assessed. Instead, we explored the relationship between the Shannon index of tree and shrub species within the 4SFI.

### 2.3. Harvesting intensity, forest structure, disturbances and climate

We defined harvesting at the plot level when at least one adult tree alive in the first census (i.e. 2SFI in the 23SFI period and 3SFI in the 34SFI period) was absent in the second census. Therefore, harvesting includes thinning, clearcut, selective logging, salvage logging after disturbances and any other harvesting type implying tree removal. With the SFI information, we calculated harvesting intensity as the percentage of basal area removed between censuses (i.e. basal area absent in 3SFI relative to 2SFI for the 23SFI period, and in 4SFI relative to 3SFI for the 34SFI period; %). To account for stand development variability, we characterised forest structure using three metrics: stand basal area, calculated as the sum of the basal area of adult trees (m^2^ ha^-1^); stand density, defined as the number of adult trees per hectare (No. trees ha^-1^); and mean tree size, representing the average d.b.h. of adult trees within each plot (mm). We used the structural variables of the initial census to relate stand development to indicator trends (i.e. 2SFI for the 23SFI period and 3SFI for the 34SFI period). To control for the occurrence of disturbances, we characterise them as the presence of at least one adult tree with signs of medium or high damage, including biotic and abiotic agents as characterised in the SFI information. To characterise spatial variations in climate, we used the CHELSA aridity index (Karger et al., 2021). The layer had a spatial resolution of 1 km^2^, which allowed us to align with the spatial resolution of the SFI. Aridity index is calculated as the ratio of the mean annual precipitation to the mean annual potential evapotranspiration over the period 1981–2010, with higher values representing wetter conditions.

### 2.4. Forest classification

To assess the impact of harvesting on indicator trends, we compared different forest types with contrasting characteristics that may influence harvesting regimes and its effect on the evaluated indicators. Forest types were classified according to tree species diversity (monospecific or mixed), protection status (protected or unprotected), stand origin (natural vs planted), and biogeographical region (Mediterranean vs temperate; see Fig. S1.1; Table S1.3). We considered a plot as monospecific if, at the initial census, a single species accounted for more than 80% of the basal area, otherwise the plot was classified as mixed (Bravo-Oviedo et al., 2014). Protected areas were characterised according to the figures of the Spanish Law on Natural Heritage and Biodiversity. Specifically, we considered national and natural parks, natural reserves and natural monuments, as these figures usually have harvesting restrictions, using SFI coordinates and the protected areas map from MITECO (https://www.miteco.gob.es/es/cartografia-y-sig/ide/descargas/biodiversidad/enp.html). We classified a plot as planted if there was evidence of soil preparation prior to planting, or if the origin of any of the three dominant tree species was artificial as recorded at the initial SFI census. Lastly, we classified plots into Mediterranean and temperate regions based on the ecoregion classification in Olson et al. (2001).

### 2.5. Statistical analyses

To assess differences in indicators over consecutive SFI censuses, we estimated marginal means for each indicator and conducted pairwise comparisons using Tukey’s test with multiple comparisons adjustment with the *emmeans* R package (Lenth, 2017). To explore the effects of harvesting on indicator trends depending on the forest type, we performed a two-step approach to assess the patterns of tree harvesting occurrence and, when tree harvest occurs, evaluate the effects of harvesting intensity.

As a first step, we fitted separate models for the trends in each of the five indicators (linear mixed models for aboveground carbon stocks, structural diversity, and the tree Shannon index, and linear models for the dominance of native tree species and standing deadwood) within each of the four forest classification (i.e. depending on diversity, protection status, origin and region), resulting in a total of twenty models. For the carbon stocks analyses, we focused on aboveground carbon stocks, as tree harvesting does not significantly affect belowground biomass in comparison to aboveground and harvesting data in the SFI account exclusively for the aerial portion of trees removed. Prior to the analyses, trends in aboveground carbon stocks were transformed using a signed square root (i.e. square root transformation preserving the sign of the original value). Trends in Shannon index and standing deadwood were rescaled to the unit interval [0, 1] using min-max normalisation and subsequently adjusted to the open interval (0, 1) to meet beta regression assumptions (Smithson & Verkuilen, 2006), and modelled using a logit link function. Plot identity was used as a random effect for the carbon stocks, structural diversity and tree Shannon index models to adjust for the temporal dependency resulting from repeated censuses over the same plots (Zuur et al., 2009). For the dominance analysis, we excluded the plots where no changes were observed because they corresponded to plots where the dominance was 100%, and the indicator could not increase further, resulting in no repeated measures over same plots.

In each first step models, we used as fixed terms initial stand structure (i.e. stand basal area, tree density and tree size), the initial value of the target indicator (i.e. initial aboveground carbon stocks, structural diversity, tree species diversity, dominance of native tree species and standing deadwood), the aridity index, the occurrence of disturbances between the SFI periods, two adjust variables (i.e. No. years between censuses to relativise the trends in indicators and the SFI period to adjust the timing of the measurement intervals), and the interaction between the forest classification (with two factor levels in each) and harvesting occurrence. The continuous fixed terms were standardized (Schielzeth, 2010) and we checked their correlations to interpret the relative contribution of the variables (Fig. S3.1). Initial basal area and initial aboveground carbon stocks were positively correlated and consequently stand basal area was not included in the carbon trend models. To assess spatial autocorrelation in the data, we used a Monte-Carlo permutation test (Manly, 2006) for Moran’s I statistic with the *spdep* R package (Bivand, 2005; see Table S3.1).

As a second step, we selected plots with harvesting occurrence and tested the effect of harvesting intensity. The reduced number of observations with tree harvesting (c. 75% of the original data, see Table S3.2) decreased the number of repeated measures over same plots in the first and second period. Therefore, for plots repeated in both censuses (∼10% of observations), we calculated the mean value for the trend indicators and fixed terms. For each forest classification (depending on diversity, protection, origin and region) and indicator (aboveground carbon stocks, structural diversity, and tree species diversity) we fitted separate linear models resulting in a total of twelve models. Each model tested the effect of harvesting intensity on the corresponding indicator within the specific forest classification. The fixed terms included the initial stand structure (i.e. stand basal area, tree density and tree size), initial indicator state for each model (i.e. aboveground carbon stocks, structural diversity and tree species diversity), climate (i.e. aridity index), No. years between censuses, and the interaction between each forest classification and harvesting intensity. Harvesting intensity was included in the model as a quadratic term to account for potential non-linear relationships between indicator trends and harvesting. Standing deadwood and the dominance of native species could not be assessed due to insufficient number of plots with harvest across forest types. We used *glmmTMB* R package to fit the models (Brooks et al., 2017) and we obtained R^2^ for the linear models and marginal and conditional R^2^ for the linear mixed models (i.e. for the fixed and fixed plus random effects, respectively; Nakagawa & Schielzeth, 2013). Finally, we diagnosed the distribution of the residuals for each fitted model (see Fig. S3.2 and Fig. S3.3). All calculations were done using in R 4.4.1 (R Core Team, 2025).

## 3. RESULTS

### 3.1. Trends in forest condition indicators

Forest condition indicators significantly increased over time in continental forests of Spain regardless of the forest type (Fig. S4.1), excepting for the dominance of native species that decreased in the first period, but remained constant in the second (Fig. 1; Table S4.1). The increases in aboveground carbon stocks per year were larger in the second analysed period (i.e. between 3SFI and 4SFI; see Fig. S4.2 and Table S4.2). The highest increases from 2SFI to 4SFI occurred in belowground and aboveground carbon stocks (i.e. mean increases of 56.1% and 62.7%, respectively), structural diversity (i.e. mean increase of 10.0%) and Shannon index (i.e. mean increase of 38.9%; Fig. 1a – c; Table S4.1). Despite this increase in Shannon diversity, it is worth noting that shrub diversity was twice as high as tree diversity in Spanish forests; however, these values cannot be compared between SFI censuses (Fig. 1c). The dominance of native species showed a small but statistically significant decrease (i.e. -0.63%; Fig. S4.3). Standing deadwood significantly increased by 83.8% from the 3SFI to 4SFI (Fig. 1d; Table S4.1).

**Figure 1.**
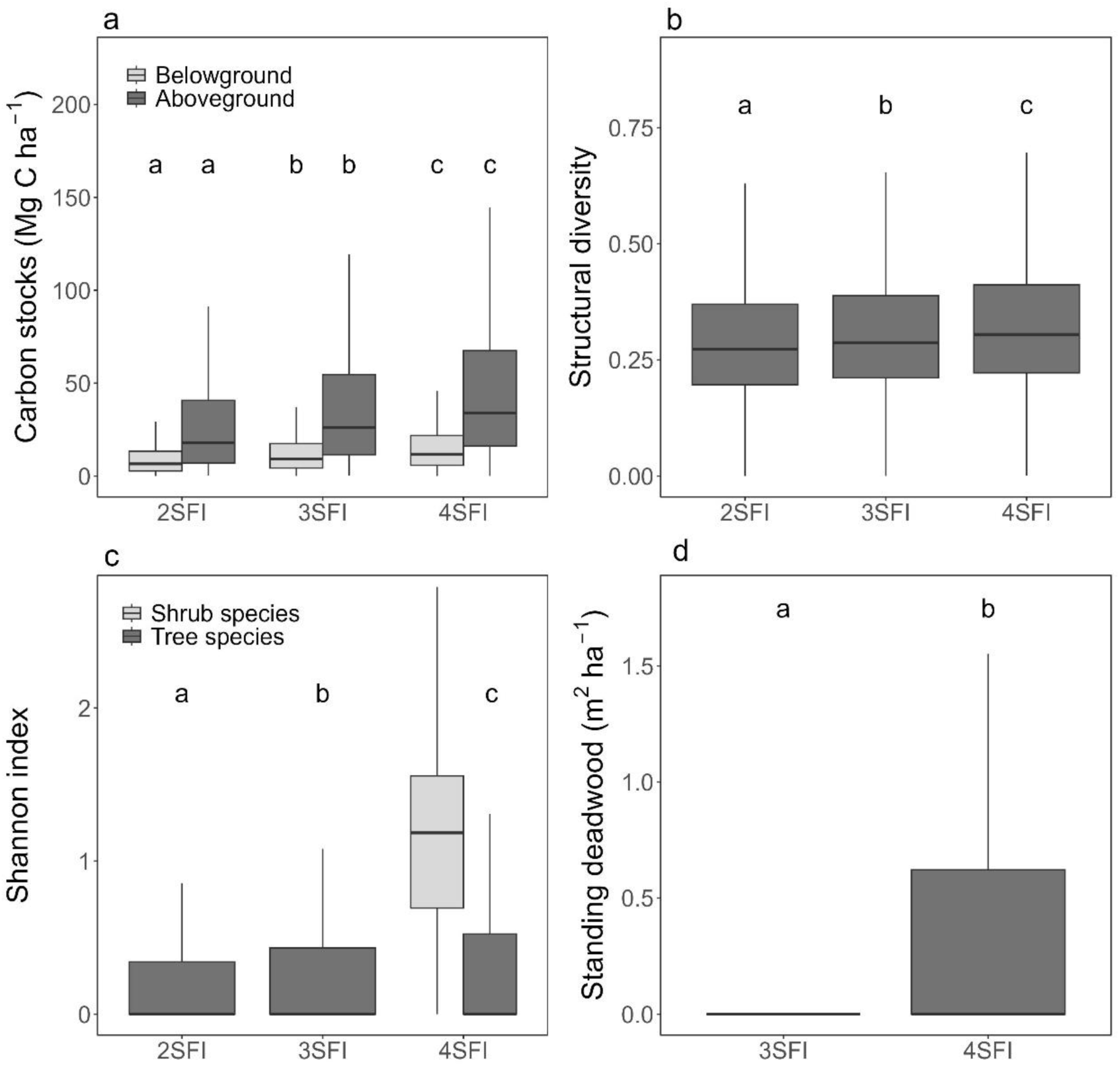
Boxplots of (a) aboveground and belowground carbon stocks (Mg C ha^-1^); (b) structural diversity; (c) Shannon index; and (d) standing deadwood (m^2^ ha^-1^) across the three consecutive Spanish Forest Inventory censuses (i.e. for the second, third and fourth: 2SFI, 3SFI and 4SFI). Inset letters represent groups with significant differences (p < 0.05) between each census. Outliers are not shown to enhance visual representation.

### 3.2. Effects of harvesting occurrence on forest condition indicators

Tree harvesting occurrence had significant effects on aboveground carbon stocks, structural diversity, Shannon index and standing deadwood trends regardless of the forest type, except for Shannon index in natural stands and within protected areas. In contrast, harvesting only affected significantly the dominance of native species in monospecific stands (Fig. 2; Table S4.3). Overall, the increases in the indicators were smaller or even decreased where harvesting occurred (Fig. 2, Table S4.3). With harvesting occurrence, the increases in aboveground carbon stocks and tree Shannon index were smaller than in unharvested stands, being close to zero for carbon stocks (except negative values for carbon stocks in temperate forests and Shannon index in mixed forests, see Fig. 2a, c). Harvesting led to negative trends in structural diversity and standing deadwood (i.e. see negative value in Fig. 2b, e), except for structural diversity in mixed forests and the temperate region, where the trend was less positive compared to unharvested plots. Additionally, decreases in the dominance of native species were smaller with harvesting occurrence in monospecific stands (Fig. 2d, Table S4.3).

**Figure 2.**
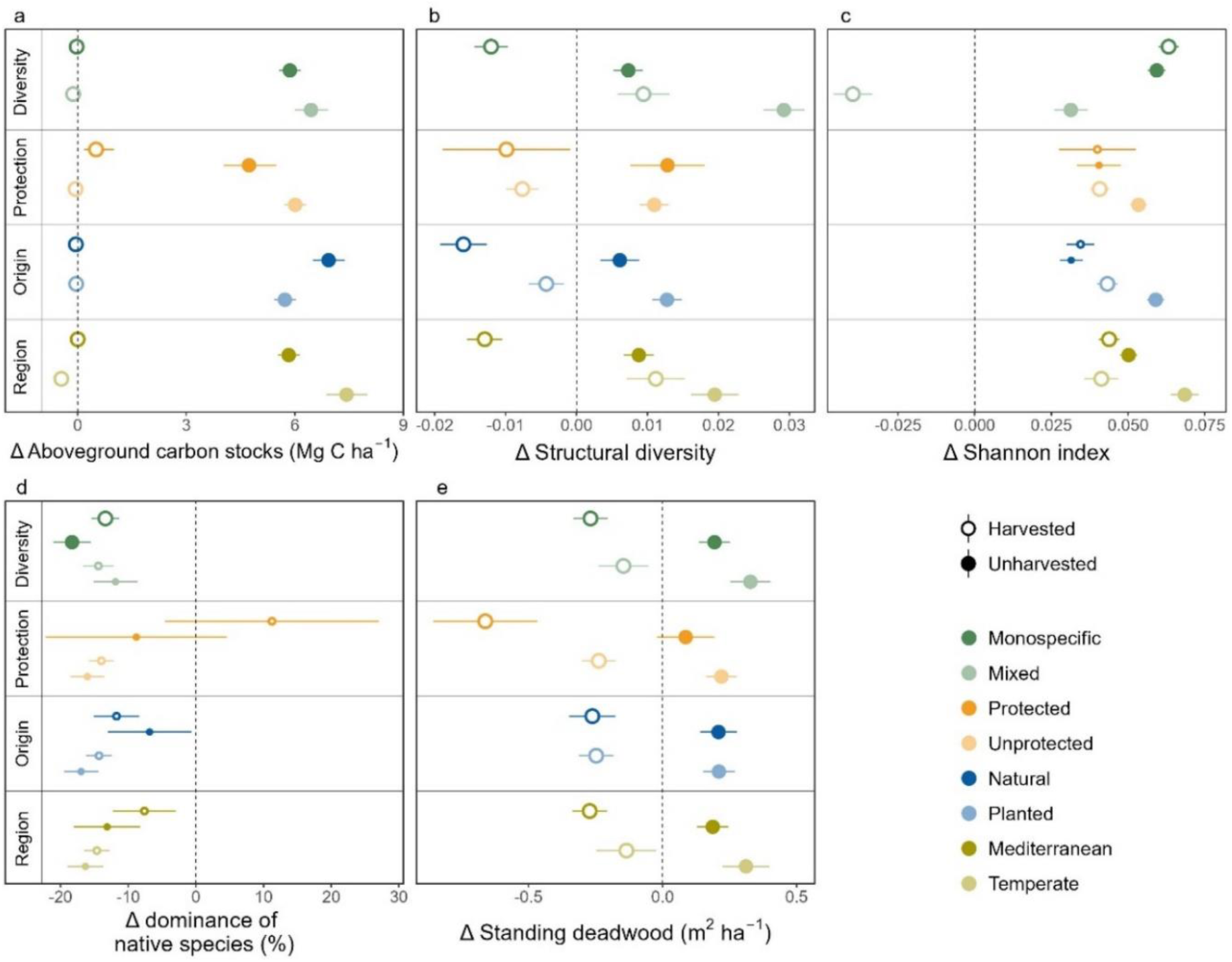
Estimate and 95% confidence intervals of trends in (a) aboveground carbon stocks (Mg C ha^-1^), (b) structural diversity, (c) Shannon index, (d) dominance of native species (%), and (e) standing deadwood (m^2^ ha^-1^) for each forest classification depending on harvesting occurrence. Small points indicate non-significant differences depending on harvesting in the same forest type (pairwise post-hoc test, p > 0.05).

### 3.3. Effects of harvesting intensity on forest condition indicators

The effects of harvesting intensity on indicator trends varied depending on the specific indicator (Fig. 3). Aboveground carbon stocks significantly decreased over time as harvesting intensity increased for all the analysed forest types, with changes in aboveground carbon stocks from positive to negative at harvesting intensities between 30% and 50% (Fig. 3a – d; Table S4.4). The greatest differences in aboveground carbon stocks between forest types were found when comparing the lowest and the highest harvesting intensities between regions, with the Mediterranean region having smaller decreases than the temperate region (Fig. 3d). Trends in structural diversity along harvesting intensity varied across the studied forest types (Fig. 3e – h), being independent of harvesting intensity in protected areas and in the temperate region (Fig. 3f – h; Table S4.4). Structural diversity showed its greatest decrease at medium harvesting intensities except for mixed stands, where it increased even with high harvesting intensities, albeit at a slower rate than under low harvesting intensity (Fig. 3e). Trends in Shannon index also varied with harvesting intensity depending on the forest types (Fig. 3i – l), but they were not significant in protected areas, natural stands and the Mediterranean region (Fig. 3j – l; Table S4.3). In unprotected areas, planted stands and the temperate region, medium harvesting intensities were related to maximum positive changes in the Shannon index (Fig. 3j – l). Trends in tree species diversity clearly differed between monospecific and mixed stands: in monospecific stands, they increased with harvesting intensity, whereas in mixed stands declined as harvesting intensity increased (Fig. 3i).

**Figure 3.**
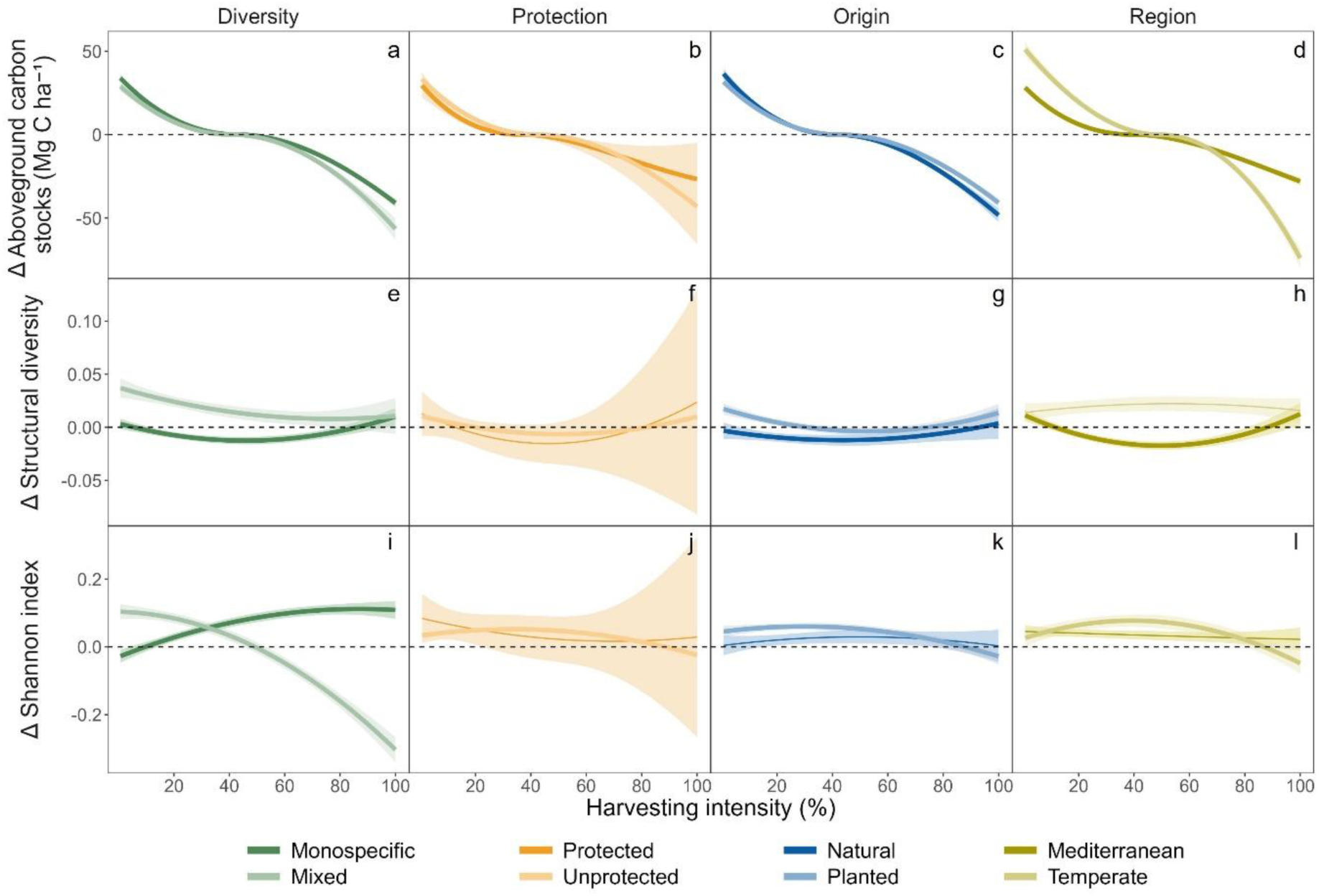
Predictions of trends in (a) aboveground carbon stocks (Mg C ha^-1^), (b) structural diversity and (c) Shannon index for each forest classification within the range of existing harvesting intensity (basal area removed between consecutive censuses; %). Wider lines represent significant effect for the forest type. Filled area represents 95% confidence intervals.

## 4. DISCUSSION

We provide a large-scale application of forest condition indicators included in the EU Nature Restoration Regulation (NRR) using National Forest Inventory data. We revealed an overall increasing trend in indicators (aboveground and belowground carbon stocks, structural diversity, Shannon index and standing deadwood) in Spanish forests from 1986 to 2022 (i.e. 56.1%; 62.7%; 10.0%; 38.9%, and 83.8%; respectively), with the exception of the dominance of native species. In addition, we evaluated the contribution of harvesting to indicator trends, with the occurrence of harvesting significantly decreasing aboveground carbon stocks, structural diversity, Shannon index and standing deadwood trends compared to unharvested plots. Harvesting intensity modulated indicator trends, being dependent on the target indicator and forest type. Altogether, our results provide evidence on how forest management influences forest condition across contrasting forest landscapes in the Iberian Peninsula, highlighting the importance of matching management practices to restoration goals under the NRR.

### 4.1. Increasing trends in forest condition indicators

All analysed indicators, except the dominance of native species increased over time in Iberian forests. Increases in carbon stocks are consistent with patterns observed in previous studies in Spanish forests; although the increases have been smaller in recent decades (Astigarraga et al., 2020; Tijerín-Triviño et al., 2025; Vayreda et al., 2012). These increases could be partially driven by the aboveground and belowground biomass accumulation underlaid by the abandonment of forest management and regrowth of secondary forests on formerly cultivated fields (Cruz-Alonso et al., 2019; Vadell et al., 2022). Furthermore, the increasing structural and taxonomic diversity may be driven by forest succession, as Iberian forests are relatively young and in early successional stages (Ruiz-Benito et al., 2014) and tree species can be gradually incorporated into secondary forests regenerating after agricultural abandonment (Espelta et al., 2020; Sánchez de Dios et al., 2023). Similarly, the amount of standing deadwood increases as the forest matures (Sandström et al., 2007). The dominance of native species was the only indicator slightly decreasing over time. This pattern aligns with studies reporting increased establishment and spread of non-native species across the Iberian Peninsula (Lara-Romero et al., 2022). Such trends may reflect the legacy of private ownership policies which prioritise the planting of fast-growing species such as *Pinus radiata* and *Eucalyptus spp.* for timber production across different regions in Spain (Rodríguez, 2006; Xunta de Galicia, 2021). The trend of dominance of native species was only positive in protected areas with harvesting actions, maybe reflecting the efforts for removing non-native species in areas designated for conservation. This suggest that restoration policies should focus on expanding the proportion of Iberian forests dominated by native species, as highlighted by the NRR.

### 4.2. Harvesting as a key factor determining temporal trends of forest condition indicators

Harvesting occurrence strongly decreased the magnitude or even reversed the direction of the studied indicator trends. Specifically, when harvesting occurred, changes over time approached zero for carbon stocks, showed smaller increases for the Shannon index and decreases for structural diversity and standing deadwood. Additionally, harvesting intensity modulated indicator trends over time, with effects varying depending on the indicator and forest type.

The decline in aboveground carbon stocks under harvesting occurrence likely reflects that, although harvesting can stimulate growth in remaining trees, as reflected by increases in carbon stocks at low intensity harvesting in our results, this increase does not offset the biomass losses from removed trees, ultimately reducing overall carbon stocks in forests (Powers et al., 2011). Regarding carbon compartments in forests other than trees, some studies have reported that harvesting may enhance resource availability such as light, water and nutrients for understory vegetation, promoting carbon accumulation in the understory (Zhou et al., 2013). In contrast, other studies have reported a limited effect on soil organic carbon (the original indicator in the NRR) than in biomass, attributed to reduced litter inputs and enhanced decomposition rates under altered soil environment (Mayer et al., 2020).

When considering aboveground carbon dynamics, we observed that high harvesting intensity resulted in greater carbon losses in the temperate than in the Mediterranean region. This likely reflects the higher overall biomass accumulation and removal in temperate forests, supported by higher water availability and higher growth rates (Ruiz-Benito et al., 2014). In addition, Mediterranean forests may be better adapted to relative intense harvesting, as their fire-adapted species allows for rapid resprouting and regeneration under increased light availability (Moghli et al., 2022).

The decreased structural diversity with harvesting occurrence could be attributed to the productive objectives of harvesting, which prioritise fast-growing and resprouting species (i.e. *Populus* and *Eucalyptus*) intensively managed for biomass production. These stands are typically subject to clearcutting or coppicing under short rotation cycles, that produce even-aged stands (Vadell et al., 2022). Our results indicate a slight increase in structural diversity in some forest types under low harvesting intensities. We anticipated a positive response at low intensities because such interventions can create canopy gaps that favour regeneration and promote heterogeneity among age classes (Pretzsch & Hilmers, 2024), particularly when combined with longer rotation periods that further enhance structural diversity (Barbeito et al., 2009). However, as harvesting intensity increases, the observed patterns diverge from our expectations: structural diversity changes were negative at medium harvesting intensity and remained close to zero at higher harvesting intensities. Previous studies suggest negative effects under very intense harvesting (Storch et al., 2019), probably associated with regeneration cuts of fast-growing stands (Torras et al., 2012; Vadell et al., 2022).

Harvesting occurrence also led to decreases in Shannon index over time, with intermediate harvesting intensities leading to the highest increases in taxonomic diversity. Increases in tree diversity in unharvested stands are consistent with other studies (Sánchez de Dios et al., 2023), highlighting the ongoing successional forests dynamics (Pecl et al., 2017), while increases in harvested stands might be related to thinning and other forest management interventions that can increase species richness according to the intermediate disturbance hypothesis (Connell, 1978; Li et al., 2014; Wu et al., 2018). Moderate disturbances can create niches and facilitate the coexistence of pioneer and late successional species, increasing tree diversity (Battles et al., 2001; Schumann et al., 2003; Zhu et al., 2007). However, the effects of harvesting on tree diversity can strongly depend on the objectives of the management interventions. Productive approaches, including clearcutting and shelterwood methods, have been associated with reductions in tree species diversity (Torras et al., 2012). In this sense, the contrasting patterns between tree species diversity and structural diversity across harvesting intensities suggest a potential trade-off. Medium harvesting intensities may promote tree species diversity by enhancing recruitment opportunities and reducing competitive exclusion (Ricklefs, 1977; Van Der Meer et al., 1999), yet the newly established individuals could often belong to the same age cohort and share similar sizes, resulting in reduced structural diversity. In this sense, thinning, and selection cutting with intermediate harvesting intensity are associated with species diversity enhancement in the Iberian Peninsula (Torras et al., 2012). However, owing to limitations in the available data, it was not possible to explicitly evaluate the effects of different harvesting types. Further research is therefore needed to better monitor the impact of different harvesting interventions on forest conditions indicators and, thus, for management recommendation for biodiversity enhancement. The observed decrease in tree species diversity with increased harvesting intensity in mixed stands may be attributed to the limited capacity of already mixed forests in the Iberian Peninsula to accommodate additional species. This limitation arises from the limited availability of functionally distinct species, as most of Spanish forests are dominated by *Pinus* and *Quercus* species (Blondel & Aronson, 1999).

The reduction of standing deadwood with harvesting occurrence may suggest that harvesting implies the removal of dead and declining trees (Gibb et al., 2005; Powers et al., 2011). Deadwood in its different forms (snags, standing and lying deadwood) is a key structural component of forest ecosystems, contributing to forest biodiversity conservation and the carbon cycle (Harmon et al., 1986; Teodosiu & Bouriud, 2012). Overall, our study highlights the value of long-term, large-scale datasets for assessing forest condition indicators. Forest inventories offer an opportunity to monitor natural dynamics and the effects of management interventions on a wide range of indicators and conditions. Integrating forest inventory data with restoration objectives can help identify management practices that enhance ecosystem services while minimizing negative impacts, such as reduced structural diversity or carbon stock losses.

### 4.3. Addressing data gaps: forest inventories and the challenges for the EU Restoration Regulation indicators

Despite the advantages of the long-term and large-scale data provided by forest inventories, we acknowledge some limitations relevant to our study. First, there is incomplete information on restoration actions implemented within the surveyed plots, which may be key for translating inventory data into practical applications. Although we could identify planted forests, they are biased to those that already have adult trees, missing the indicators state at initial stages after planting. Also, planted forests might include a variety of situations from old, naturalised plantations to productive short-rotation stands.

Some proposed indicators by the NRR (i.e. lying deadwood, forest connectivity) could not be assessed due to the absence of specific data. Furthermore, forest inventories are primarily designed to measure aboveground components and thus provide limited capacity to assess soil conditions or to infer relationships between harvesting and soil carbon sequestration. These data gaps highlight the potential to complement forest inventory information with additional sources, such as high-resolution aerial imagery, forest maps and strategic soil sampling. Despite certain missing information, forest inventories allow the characterization of further interesting attributes related to forest condition, such as shrub diversity, since shrubs in the understory contain most of woody species diversity in Mediterranean forests (Martín-Queller et al., 2011), and tree regeneration on different stages, strongly linked to natural tree resilience. Those indicators can be integrated to estimate the success of restoration efforts (Cruz-Alonso et al., 2019).

We also encountered other challenges related to the proposed indicators of the NRR. For example, the indicator *share of forest dominated by native species* may have limited applicability depending on the spatial scale considered. In the case of SFI data (i.e. 1 km^2^ grid cells), nearly 95% of the plots were already fully dominated by native species. Since the NRR requires an increase in this indicator, achieving such a target when the indicator is already at its maximum would be unfeasible and meaningless for most Spanish forests, even those that are degraded but 100% composed by native species. Similarly, interpreting an increase in standing deadwood as a sign of restoration success might be confusing, as it may instead reflect higher tree mortality rather than successful restoration actions.

## 5. CONCLUSIONS

Forest condition indicators (i.e. aboveground and belowground carbon stocks, structural diversity, tree species diversity and standing deadwood) increased over time, with the most pronounced gains occurring in recent decades. Our results show reductions in the trends of the evaluated indicators associated with harvesting occurrence across all forest types; with trends becoming negative for aboveground carbon stocks, structural diversity and standing deadwood. Our findings also revealed contrasting effects of harvesting intensity on forest condition indicators. Aboveground carbon stocks increased under low harvesting intensity but declined at medium to high intensities across forest types. Structural diversity decreased at intermediate harvesting intensity while trends in tree species diversity increased at intermediate harvesting intensity.

Our results suggest that future harvesting practices can be tailored to the promotion of carbon sequestration, uneven-aged stand structures, tree diversity, and greater amounts of deadwood thereby enhancing indicators and supporting the objectives of the NRR. Maximizing all these outcomes simultaneously may be infeasible at the stand level and should be planned at a landscape level. Importantly, management decisions need to account for the response of multiple indicators, as focusing on a single variable may overlook important trade-offs. Low-intensity harvesting can enhance the indicators and support the objectives of the NRR while using timber resources and reducing vulnerability to disturbances; yet outcomes vary with forest type, site conditions and silvicultural systems, requiring context-specific guidelines. In the absence of productive objectives, maintaining unharvested conditions may be equally or even more beneficial for most indicators.

Forest inventories stand out as a key tool for monitoring restoration progress across large spatial and long temporal scales. They provide a minimum level of indicator monitoring for all administrations in a country. By incorporating stand-level information, they also offer practical guidance on spatial scale, which is not explicitly provided in the NRR. The information about forest condition in forest inventories can aid in establishing reference levels for forests indicators within each EU member. Finally, the naturally positive trend of most indicators highlights the need to establish sufficiently strict thresholds of increase to evaluate active forest restoration, or minimum rates of increase that indicates acceptable conditions, as observed in natural forests.

## ACKNOWLEDGEMENTS

This study was supported by the grant FUNFOREST (3060/2023, funded by Organismo Autónomo Parques Nacionales and Ministerio para la Transición Ecológica y el Reto Demográfico) and LARGE (N° PID2021-123675OB-C41, Science and Innovation Ministry, Agencia Estatal de Investigation). PR is funded by an assistant professor grant from the Complutense University of Madrid. JA was supported by the Basque Government’s Postdoctoral Programme for the Improvement of Doctoral Research Staff (POS_2024_1_0026). VCA is co-supported by the Community of Madrid under the 2024 call for the ‘César Nombela’ research talent attraction programme (2024-T1/ECO-31335). We acknowledge the availability of open-access data from the Spanish National Inventory, and all the help provided to their use from the Ministerio para la Transición Ecológica (https://www.miteco.gob.es/es/biodiversidad/temas/inventarios-nacionales.html).

## AUTHOR CONTRIBUTIONS

PR, VCA, EA and PRB planned the research. PR, VCA, PRB, JA, MAZ and SS processed the data. VCA, EA, PRB and MAZ refined the study approach and PR, VCA and JA analysed the data and PR and VCA designed the graphical representation. All authors contributed to the final version through several revisions.

