## Supporting_information for "Harvesting effects on forest condition indicators across Iberian forests: Implications for the EU Nature Restoration Regulation"

**Appendix S1. Details of forest condition indicators calculations**

**Appendix S2. Stocks of organic carbon in litter and mineral soil according to the Spanish Forest Inventory (SFI).**

**Appendix S3. Further modelling details of changes in forest conditions indicators.**

**Appendix S4. Complementary results of changes in forest conditions indicators.**

**References**

**Appendix S1. Details of forest condition indicators calculations**

**Table S1.1.** Dry biomass equations for the tree species used in the study. Carbon content of each species was taken from Montero et al. (2005). $W_s$: Biomass weight of the stem fraction (kg); $W_{bk}$: Biomass weight of the stem bark fraction (kg); $W_{b7}$: Biomass weight of the thick branches fraction (diameter larger than 7 cm) (kg); $W_{b2\text{-}7}$: Biomass weight of medium branches fraction (diameter between 2 and 7 cm) (kg); $W_{b2}$: Biomass weight of thin branches fraction (diameter smaller than 2 cm) (kg); $W_t$: Biomass weight of twigs fraction (kg); $W_f$: Biomass weight of foliar (leaves or needles) fraction (kg); $W_r$: Biomass weight of the belowground fraction (kg); DBH.: diameter at breast height (cm); h: tree height (m).

| **Species** | **Source** | **Equation** | **Carbon content (%)** |
|---|---|---|---|
| *Abies alba* | Ruiz-Peinado et al., (2011) | $W_s = 0.0189 \cdot DBH^2 \cdot h$<br>$W_{b7}+W_{b27} = 0.0584 \cdot DBH^2$<br>$W_{b2+n} = 0.0371 \cdot DBH^2 + 0.968 \cdot h$<br>$W_r = 0.101 \cdot DBH^2$ | 50.6 |
| *Abies pinsapo* | Ruiz-Peinado et al. (2011) (Roots from *Abies alba*) | $W_s = 0.00960 \cdot DBH^2 \cdot h$<br>$W_{b7}$ (if DBH > 32.5) $= 1.637 \cdot (DBH-32.5)^2 - 0.0719 \cdot (DBH-32.5)^2 \cdot h$<br>$W_{b2\text{-}7} = 0.00344 \cdot DBH^2 \cdot h$<br>$W_{b2+n} = 0.131 \cdot DBH \cdot h$ | 50.0 |
| *Alnus glutinosa* | Ruiz-Peinado et al., (2012) | $W_s = 0.0191 \cdot DBH^2 \cdot h$<br>$W_{b7} + W_{b2\text{-}7} = 0.0512 \cdot DBH^2$<br>$W_{b2+l} = 0.0567 \cdot DBH \cdot h$<br>$W_r = 0.214 \cdot DBH^2$ | 50.0 |
| *Castanea sativa* | Ruiz-Peinado et al. (2012) | $W_s = 0.0142 \cdot DBH^2 \cdot h$<br>$W_{b7}$ (if DBH > 12.5) $= 0.223 \cdot (DBH - 12.5)^2$<br>$W_{b2\text{-}7} = 0.230 \cdot DBH \cdot h$<br>$W_{b2} = 0.221 \cdot DBH \cdot h$<br>$W_r = 0.0211 \cdot DBH^{2.804}$ | 48.4 |
| *Ceratonia siliqua* | Ruiz-Peinado et al. (2012) | $W_s = 0.142 \cdot DBH^{1.974}$<br>$W_{b7} = 0.104 \cdot DBH^2$<br>$W_{b2\text{-}7} = 0.0538 \cdot DBH^2$<br>$W_{b2+l} = 0.151 \cdot DBH^2 - 0.00740 \cdot DBH^2 \cdot h$<br>$W_r = 0.335 \cdot DBH^2$ | 50.0 |
| *Eucalyptus camaldulensis* | From *Eucalyptus globulus* | $W_{s+b7} = 0.0221 \cdot DBH^2 \cdot h$<br>$W_{b2\text{-}7} = 0.154 \cdot DBH^{1.668}$<br>$W_{b2+l} = 0.180 \cdot (DBH^2 \cdot h)^{0.587}$ | 47.5 |
| *Eucalyptus globulus* | Ruiz-Peinado et al. (2012) | $W_{s+b7} = 0.0221 \cdot DBH^2 \cdot h$<br>$W_{b2\text{-}7} = 0.154 \cdot DBH^{1.668}$<br>$W_{b2+l} = 0.180 \cdot (DBH^2 \cdot h)^{0.587}$ | 47.5 |
| *Eucalyptus gomphocephalus* | From *Eucalyptus globulus* | $W_{s+b7} = 0.0221 \cdot DBH^2 \cdot h$<br>$W_{b2\text{-}7} = 0.154 \cdot DBH^{1.668}$<br>$W_{b2+l} = 0.180 \cdot (DBH^2 \cdot h)^{0.587}$ | 47.5 |

| | | | |
|---|---|---|---|
| *Eucalyptus nintens* | From *Eucalyptus globulus* | $W_{s+b7} = 0.0221 \cdot DBH^2 \cdot h$<br>$W_{b2\text{-}7} = 0.154 \cdot DBH^{1.668}$<br>$W_{b2+l} = 0.180 \cdot (DBH^2 \cdot h)^{0.587}$ | 47.5 |
| *Eucalyptus spp.* | From *Eucalyptus globulus* | $W_{s+b7} = 0.0221 \cdot DBH^2 \cdot h$<br>$W_{b2\text{-}7} = 0.154 \cdot DBH^{1.668}$<br>$W_{b2+l} = 0.180 \cdot (DBH^2 \cdot h)^{0.587}$ | 47.5 |
| *Eucalyptus viminalis* | From *Eucalyptus globulus* | $W_{s+b7} = 0.0221 \cdot DBH^2 \cdot h$<br>$W_{b2\text{-}7} = 0.154 \cdot DBH^{1.668}$<br>$W_{b2+l} = 0.180 \cdot (DBH^2 \cdot h)^{0.587}$ | 47.5 |
| *Fagus sylvatica* | Ruiz-Peinado et al. (2012) | $W_s = 0.0676 \cdot DBH^2 + 0.0182 \cdot DBH^2 \cdot h$<br>$W_{b7}$ (if DBH > 22.5) $= 0.830 \cdot (DBH - 22.5)^2 - 0.0248 \cdot (DBH - 22.5)^2 \cdot h$<br>$W_{b2\text{-}7} = 0.0792 \cdot DBH^2$<br>$W_{b2} = 0.0930 \cdot DBH^2 - 0.00226 \cdot DBH^2 \cdot h$<br>$W_r = 0.106 \cdot DBH^2$ | 48.6 |
| *Fraxinus angustifolia* | Ruiz-Peinado et al. (2012) | $W_s = 0.0296 \cdot DBH^2 \cdot h$<br>$W_{b7}$ (if DBH > 12.5) $= 0.231 \cdot (DBH - 12.5)^2$<br>$W_{b2\text{–}7} = 0.0925 \cdot DBH^2$<br>$W_{b2} = 2.005 \cdot DBH$<br>$W_r = 0.359 \cdot DBH^2$ | 47.8 |
| *Fraxinus excelsior* | From *Fraxinus angustifolia* | $W_s = 0.0296 \cdot DBH^2 \cdot h$<br>$W_{b7}$ (if DBH > 12.5) $= 0.231 \cdot (DBH - 12.5)^2$<br>$W_{b2\text{–}7} = 0.0925 \cdot DBH^2$<br>$W_{b2} = 2.005 \cdot DBH$<br>$W_r = 0.359 \cdot DBH^2$ | 47.8 |
| *Juniperus communis* | From *Juniperus thurifera* | $W_s = 0.0132 \cdot DBH^2 \cdot h + 0.217 \cdot DBH \cdot h$<br>$W_{b7}$ (if DBH > 22.5) $= 0.107 \cdot (DBH\text{-}22.5)^2$<br>$W_{b2\text{-}7} = 0.00792 \cdot DBH^2 \cdot h$<br>$W_{b2+n} = 0.273 \cdot DBH \cdot h$<br>$W_r = 0.0767 \cdot DBH^2$ | 50.0 |
| *Juniperus oxycedrus* | From *Juniperus thurifera* | $W_s = 0.0132 \cdot DBH^2 \cdot h + 0.217 \cdot DBH \cdot h$<br>$W_{b7}$ (if DBH > 22.5) $= 0.107 \cdot (DBH\text{-}22.5)^2$<br>$W_{b2\text{-}7} = 0.00792 \cdot DBH^2 \cdot h$<br>$W_{b2+n} = 0.273 \cdot DBH \cdot h$<br>$W_r = 0.0767 \cdot DBH^2$ | 50.0 |
| *Juniperus phoenicea* | From *Juniperus thurifera* | $W_s = 0.0132 \cdot DBH^2 \cdot h + 0.217 \cdot DBH \cdot h$<br>$W_{b7}$ (if DHB > 22.5) $= 0.107 \cdot (DBH\text{-}22.5)^2$<br>$W_{b2\text{-}7} = 0.00792 \cdot DBH^2 \cdot h$<br>$W_{b2+n} = 0.273 \cdot DBH \cdot h$<br>$W_r = 0.0767 \cdot DBH^2$ | 50.0 |
| *Juniperus thurifera* | Ruiz-Peinado et al. (2011) | $W_s = 0.0132 \cdot DBH^2 \cdot h + 0.217 \cdot DBH \cdot h$ | 47.5 |

| | | | |
|---|---|---|---|
| | | $W_{b7}$ (if DBH > 22.5) = 0.107 · (DBH-22.5)$^2$<br>$W_{b2-7}$ = 0.00792 · DBH$^2$ · h<br>$W_{b2+n}$ = 0.273 · DBH · h<br>$W_r$ = 0.0767 · DBH$^2$ | |
| *Olea europaea* | Ruiz-Peinado et al. (2012) | $W_s$ = 0.0114 · DBH$^2$ · h<br>$W_{b7}$ = 0.0108 · DBH$^2$ · h<br>$W_{b2-7}$ = 1.672 · DBH<br>$W_{b2+1}$ = 0.0354 · DBH$^2$ + 1.187 · h<br>$W_r$ = 0.147 · DBH$^2$ | 47.3 |
| *Pinus halepensis* | Ruiz-Peinado et al. (2011) | $W_s$ = 0.0139 · DBH$^2$ · h<br>$W_{b7}$ (if DBH > 27.5) = 3.926 · (DBH-27.5)<br>$W_{b2-7}$ = 4.257 + 0.00506 · DBH$^2$ · h - 0.0722 DBH · h<br>$W_{b2+n}$ = 6.197 + 0.00932 · DBH$^2$ · h - 0.0686 · DBH · h<br>$W_r$ = 0.0785 · DBH$^2$ | 49.9 |
| *Pinus nigra* | Ruiz-Peinado et al. (2011) | $W_s$ = 0.0403 · DBH$^{1.838}$ · h$^{0.945}$<br>$W_{b7}$ (if DBH > 27.5) = 3.926 · (DBH-27.5)<br>$W_{b2-7}$ = 0.0521 · DBH$^2$<br>$W_{b2+n}$ = 0.0720 · DBH$^2$<br>$W_r$ = 0.0189 · DBH$^{2.445}$ | 50.9 |
| *Pinus pinaster* | Ruiz-Peinado et al. (2011) | $W_s$ = 0.0278 · DBH$^{2.115}$ · h$^{0.618}$<br>$W_{b7}$+$W_{b27}$ = 0.000381 · DBH$^{3.141}$<br>$W_{b2+n}$ = 0.0129 · DBH$^{2.320}$<br>$W_r$ = 0.00444 · DBH$^{2.804}$ | 51.1 |
| *Pinus pinea* | Ruiz-Peinado et al. (2011) | $W_s$ = 0.0224 · DBH$^{1.923}$ · h$^{1.0193}$<br>$W_{b7}$ (if DBH > 22.5) = 0.247 · (DBH-22.5)$^2$<br>$W_{b2-7}$ = 0.0525 · DBH$^2$<br>$W_{b2+n}$ = 21.927 + 0.0707 · DBH$^2$ - 2.827 · h<br>$W_r$ = 0.117 · DBH$^2$ | 50.8 |
| *Pinus radiata* | From *Pinus pinaster* | $W_s$ = 0.0278 · DBH$^{2.115}$ · h$^{0.618}$<br>$W_{b7}$+$W_{b27}$ = 0.000381 · DBH$^{3.141}$<br>$W_{b2+n}$ = 0.0129 · DBH$^{2.320}$<br>$W_r$ = 0.00444 · DBH$^{2.804}$ | 49.7 |
| *Pinus sylvestris* | Ruiz-Peinado et al. (2011) | $W_s$ = 0.0154 · DBH$^2$ · h<br>$W_{b7}$ (if DBH > 37.5) = 0.540 · (DBH-37.5)$^2$ - 0.0119 · (DBH-37.5)$^2$ · h<br>$W_{b2-7}$ = 0.0295 · DBH$^{2.742}$ · h$^{-0.899}$<br>$W_{b2+n}$ = 0.530 · DBH$^{2.199}$ · h$^{-1.153}$<br>$W_r$ = 0.130 · DBH$^2$ | 50.9 |
| *Pinus uncinata* | Ruiz-Peinado et al. (2011) | $W_s$ = 0.0203 · DBH$^2$ · h<br>$W_{b7}$+$W_{b27}$ = 0.0379 · DBH$^2$<br>$W_{b2+n}$ = 2.740·DBH - 2.641 · h<br>$W_r$ = 0.193 · DBH$^2$ | 50.9 |

| | | | |
|---|---|---|---|
| *Populus alba* | From *Populus x canadiensis* | $W_s = 0.0130 \cdot DBH^2 \cdot h$<br>$W_{b7}$ (if $DBH > 22.5$) $= 0.538 \cdot (DBH - 22.5)^2 - 0.0130 \cdot (DBH - 22.5)^2 \cdot h$<br>$W_{b2-7} = 0.0385 \cdot DBH^2$<br>$W_{b2+1} = 0.0774 \cdot DBH^2 - 0.00198 \cdot DBH^2 \cdot h$<br>$W_r = 0.122 \cdot DBH^2$ | 48.3 |
| *Populus nigra* | From *Populus x canadiensis* | $W_s = 0.0130 \cdot DBH^2 \cdot h$<br>$W_{b7}$ (if $DBH > 22.5$) $= 0.538 \cdot (DBH - 22.5)^2 - 0.0130 \cdot (DBH - 22.5)^2 \cdot h$<br>$W_{b2-7} = 0.0385 \cdot DBH^2$<br>$W_{b2+1} = 0.0774 \cdot DBH^2 - 0.00198 \cdot DBH^2 \cdot h$<br>$W_r = 0.122 \cdot DBH^2$ | 48.3 |
| *Populus tremula* | From *Populus x canadiensis* | $W_s = 0.0130 \cdot DBH^2 \cdot h$<br>$W_{b7}$ (if $DBH > 22.5$) $= 0.538 \cdot (DBH - 22.5)^2 - 0.0130 \cdot (DBH - 22.5)^2 \cdot h$<br>$W_{b2-7} = 0.0385 \cdot DBH^2$<br>$W_{b2+1} = 0.0774 \cdot DBH^2 - 0.00198 \cdot DBH^2 \cdot h$<br>$W_r = 0.122 \cdot DBH^2$ | 48.3 |
| *Populus x canadiensis* | Ruiz-Peinado et al. (2012) | $W_s = 0.0130 \cdot DBH^2 \cdot h$<br>$W_{b7}$ (if $DBH > 22.5$) $= 0.538 \cdot (DBH - 22.5)^2 - 0.0130 \cdot (DBH - 22.5)^2 \cdot h$<br>$W_{b2-7} = 0.0385 \cdot DBH^2$<br>$W_{b2+1} = 0.0774 \cdot DBH^2 - 0.00198 \cdot DBH^2 \cdot h$<br>$W_r = 0.122 \cdot DBH^2$ | 48.3 |
| *Quercus canariensis* | Ruiz-Peinado et al. (2012) | $W_s = 0.0126 \cdot DBH^2 \cdot h$<br>$W_{b7} = 0.103 \cdot DBH^2$<br>$W_{b2-7} + W_{b2} + l = 0.167 \cdot DBH \cdot h$<br>$W_r = 0.135 \cdot DBH^2$ | 48.6 |
| *Quercus faginea* | Ruiz-Peinado et al. (2012) | $W_s = 0.154 \cdot DBH^2$<br>$W_{b7} = 0.0861 \cdot DBH^2$<br>$W_{b2-7} = 0.127 \cdot DBH^2 - 0.00598 \cdot DBH^2 \cdot h$<br>$W_{b2+1} = 0.0726 \cdot DBH^2 - 0.00275 \cdot DBH^2 \cdot h$<br>$W_r = 0.169 \cdot DBH^2$ | 48.0 |
| *Quercus ilex* | Ruiz-Peinado et al. (2012) | $Ws = 0.143 \cdot DBH^2$<br>$W_{b7}$ (if $DBH > 12.5$) $= 0.0684 \cdot (DBH - 12.5)^2 \cdot h$<br>$W_{b2-7} = 0.0898 \cdot DBH^2$<br>$W_{b2+1} = 0.0824 \cdot DBH^2$<br>$W_r = 0.254 \cdot DBH^2$ | 47.5 |
| *Quercus petraea* | From *Quercus pyrenaica* | $W_s + W_{b7} = 0.0261 \cdot DBH^2 \cdot h$<br>$W_{b2-7} = -0.0260 \cdot DBH^2 + 0.536 \cdot h + 0.00538 \cdot DBH^2 \cdot h$<br>$W_{b2} = 0.898 \cdot DBH - 0.445 \cdot h$ | 48.4 |

| | | | |
|---|---|---|---|
| | | $W_r = 0.143 \cdot DBH^2$ | |
| *Quercus pubescens* | From *Quercus faginea* | $W_s = 0.154 \cdot DBH^2$<br>$W_{b7} = 0.0861 \cdot DBH^2$<br>$W_{b2\text{-}7} = 0.127 \cdot DBH^2 - 0.00598 \cdot DBH^2 \cdot h$<br>$W_{b2+l} = 0.0726 \cdot DBH^2 - 0.00275 \cdot DBH^2 \cdot h$<br>$W_r = 0.169 \cdot DBH^2$ | 48.0 |
| *Quercus pyrenaica* | Ruiz-Peinado et al. (2012) | $W_s + W_{b7} = 0.0261 \cdot DBH^2 \cdot h$<br>$W_{b2\text{-}7} = -0.0260 \cdot DBH^2 + 0.536 \cdot h + 0.00538 \cdot DBH^2 \cdot h$<br>$W_{b2} = 0.898 \cdot DBH - 0.445 \cdot h$<br>$W_r = 0.143 \cdot DBH^2$ | 47.5 |
| *Quercus robur* | From *Quercus canariensis* | $W_s = 0.0126 \cdot DBH^2 \cdot h$<br>$W_{b7} = 0.103 \cdot DBH^2$<br>$W_{b2\text{-}7} + W_{b2} + l = 0.167 \cdot DBH \cdot h$<br>$W_r = 0.135 \cdot DBH^2$ | 48.6 |
| *Quercus rubra* | From *Quercus ilex* | $Ws = 0.143 \cdot DBH^2$<br>$W_{b7}$ (if DBH > 12.5) $= 0.0684 \cdot (DBH - 12.5)^2 \cdot h$<br>$W_{b2\text{-}7} = 0.0898 \cdot DBH^2$<br>$W_{b2+l} = 0.0824 \cdot DBH^2$<br>$W_r = 0.254 \cdot DBH^2$ | 47.5 |
| *Quercus suber* | Ruiz-Peinado et al. (2012) | $W_s = 0.00525 \cdot DBH^2 \cdot h + 0.278 \cdot DBH \cdot h$<br>$W_{b7} = 0.0135 \cdot DBH^2 \cdot h$<br>$W_{b2\text{–}7} = 0.127 \cdot DBH \cdot h$<br>$W_{b2+l} = 0.0463 \cdot DBH \cdot h$<br>$W_r = 0.0829 \cdot DBH^2$ | 47.2 |

**Table S1.2.** List of non-native species and family

| Species | Family |
|---|---|
| *Acacia dealbata* | Leguminosae |
| *Acacia melanoxylon* | Leguminosae |
| *Acacia spp.* | Leguminosae |
| *Acer negundo* | Aceraceae |
| *Ailanthus altissima* | Simaroubaceae |
| *Apollonias barbujana* | Lauraceae |
| *Cedrus atlantica* | Pinaceae |
| *Cedrus deodara* | Pinaceae |
| *Cedrus libani* | Pinaceae |
| *Chamaecyparis lawsoniana* | Cupressaceae |
| *Crataegus laciniata* | Rosaceae |
| *Cupressus arizonica* | Cupressaceae |
| *Cupressus lusitanica* | Cupressaceae |
| *Cupressus macrocarpa* | Cupressaceae |
| *Eucalyptus camaldulensis* | Myrtaceae |
| *Eucalyptus globulus* | Myrtaceae |
| *Eucalyptus gomphocephalus* | Myrtaceae |
| *Eucalyptus nitens* | Myrtaceae |
| *Eucalyptus viminalis* | Myrtaceae |
| *Gleditsia triacanthos* | Leguminosae |
| *Larix decidua* | Pinaceae |
| *Larix spp.* | Pinaceae |
| *Platanus hispanica* | Platanaceae |
| *Populus x canadensis* | Salicaceae |
| *Pseudotsuga menziesii* | Pinaceae |
| *Quercus rubra* | Fagaceae |
| *Robinia pseudacacia* | Salicaceae |
| *Salix babylonica* | Salicaceae |
| *Ulmus pumila* | Ulmaceae |

**Table S1.3**. Number of observations for each forest type classified by harvesting occurrence.

| | Diversity | | Origin | | Protection | | Region | |
|---|---|---|---|---|---|---|---|---|
| | Monospecific | Mixed | Natural | Planted | Protected | Unprotected | Mediterranean | Temperate |
| Harvested | 14,524 | 4,133 | 5,495 | 13,162 | 533 | 46,210 | 13,282 | 5,375 |
| Unharvested | 39,283 | 8,759 | 10,169 | 37,863 | 1,832 | 18,124 | 39,375 | 8,667 |

**Figure S1.1.** Distribution maps of the forest types classified in the study area, depending on diversity (monospecific or mixed), protection level (protected or unprotected), planted status (natural forests or planted) and biogeographical region (Mediterranean or temperate).

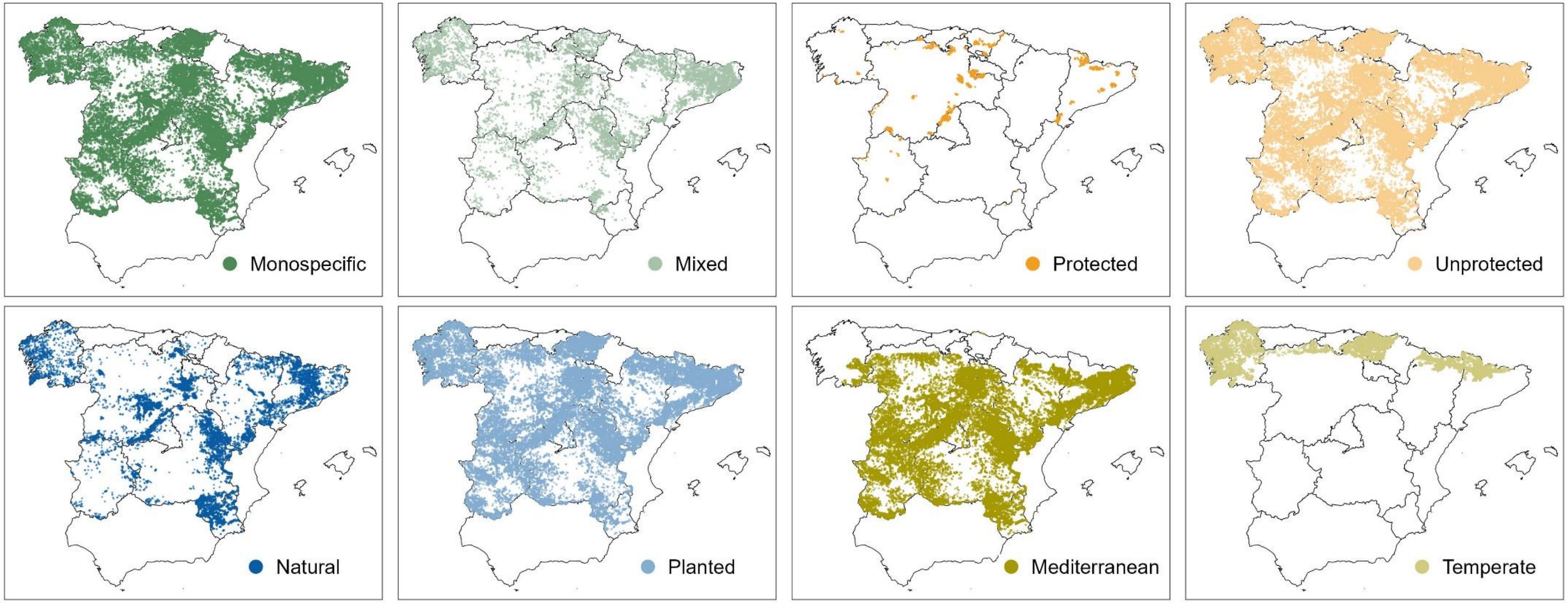

### Appendix S2. Stocks of organic carbon in litter and mineral soil according to the Spanish Forest Inventory (SFI).

Two variables measured in the SFI can be related to the stocks of organic carbon in litter and mineral soil:

#### Litter depth

In the Spanish Forest Inventory, litter depth is characterised as a coded variable according to layer thickness. A thickness of less than 0.5 cm is coded as 0, from 0.5 to 1.4 cm as 1, from 1.5 to 2.4 cm as 2, from 2.5 to 3.4 cm as 3, and so on. When plots contained areas with different litter layer thicknesses, the estimated average category was recorded.

#### Organic matter content

The organic matter content of the soil in each plot is classified according to its degree of humification. The soil is considered humus-rich soil when, at a depth of 15 cm, the purity ("value" in MUNSELL scale of soil colour) was lower than 4, or when the litter layer exceeded 5 cm in thickness and the purity at 15 cm depth was lower than 6 (MITECO, 2014). The soil is classified as moderately humus-rich soil when the purity at 15 cm depth was lower than 6 with and there is a null or very thin litter layer, or when the litter layer was thicker than 5 cm and the purity at 15 cm depth was equal to or greater than 6. In all other cases, the soil is considered humus-poor soil. Comparable data of organic matter content is available for the third and fourth Spanish Forest Inventory.

**Figure S2.1.** (a) Litter depth category according to the Spanish Forest Inventory coding system for the second, third, and fourth censuses. (b) Percentage of plots with poor, moderately and high humus for the third (3SFI) and fourth (4SFI) censuses.

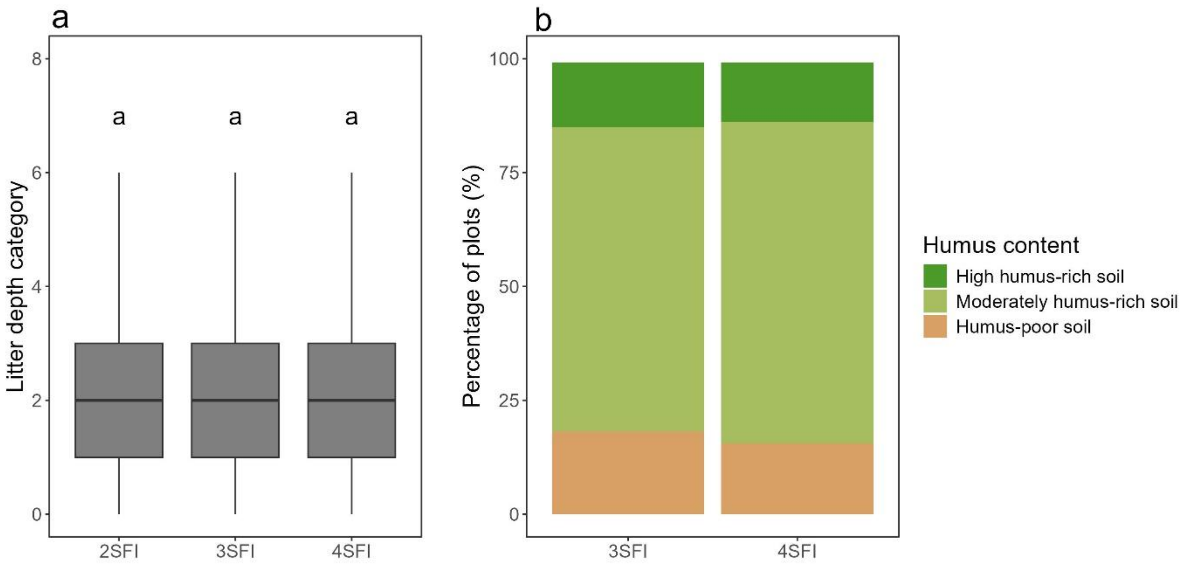

**Appendix S3. Further modelling details of changes in forest conditions indicators.**

**Table S3.1.** Results of Monte Carlo simulations (n = 999 permutations) of Moran's I for spatial autocorrelation for the residuals of each forest biodiversity indicator model. P values higher than 0.05 indicate no spatial autocorrelation.

| | **Observed statistic (Moran's I)** | **p-value** |
|---|---|---|
| Aboveground carbon storage | -0.000015 | 0.39 |
| Structural diversity | -0.000016 | 0.67 |
| Shannon Index | -0.000035 | 0.88 |
| Dominance of native species | -0.00028 | 0.09 |
| Standing deadwood | -0.000049 | 0.53 |

**Table S3.2**. Number of observations used for the statistical analyses for each indicator

| | **Aboveground carbon storage (Mg C ha$^{-1}$)** | **Structural diversity** | **Shannon Index** | **Dominance of native species (%)** | **Standing deadwood (m$^2$ ha$^{-1}$)** |
|---|---|---|---|---|---|
| Harvesting occurrence | 66,699 | 60,447 | 65,053 | 3,482 | 20,060 |
| Harvesting intensity | 16,019 | 14,547 | 16,019 | 1,893 | 1,538 |

**Figure S3.1**. Relationships between the explanatory variables for the analyses of biodiversity indicators.

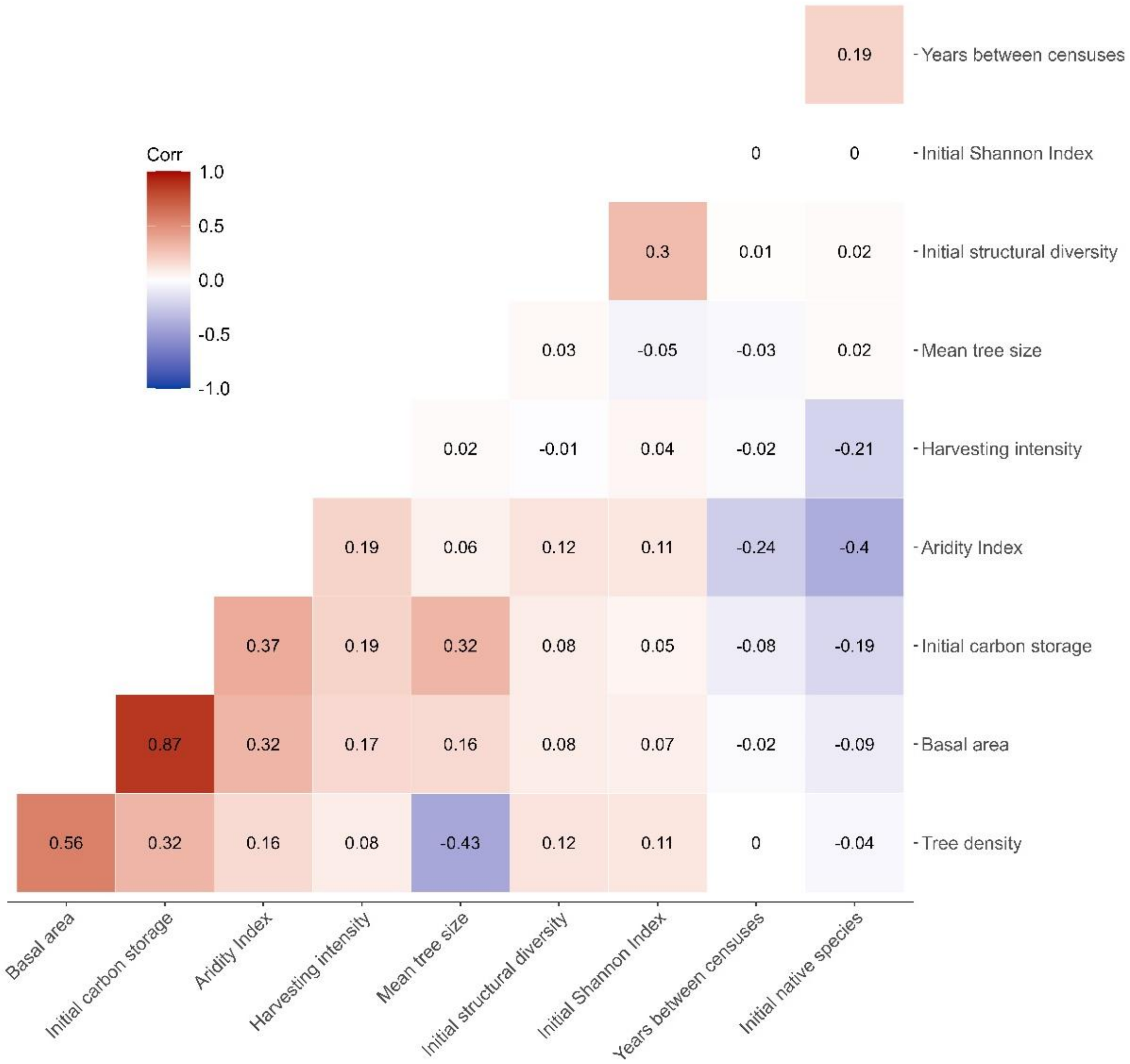

**Figure S3.2.** Scatterplot of residuals versus predicted values and histogram of the residuals resulted from mixed lineal models for changes in carbon storage, structural diversity and Shannon Index depending on the forest type and the presence or absence of harvesting.

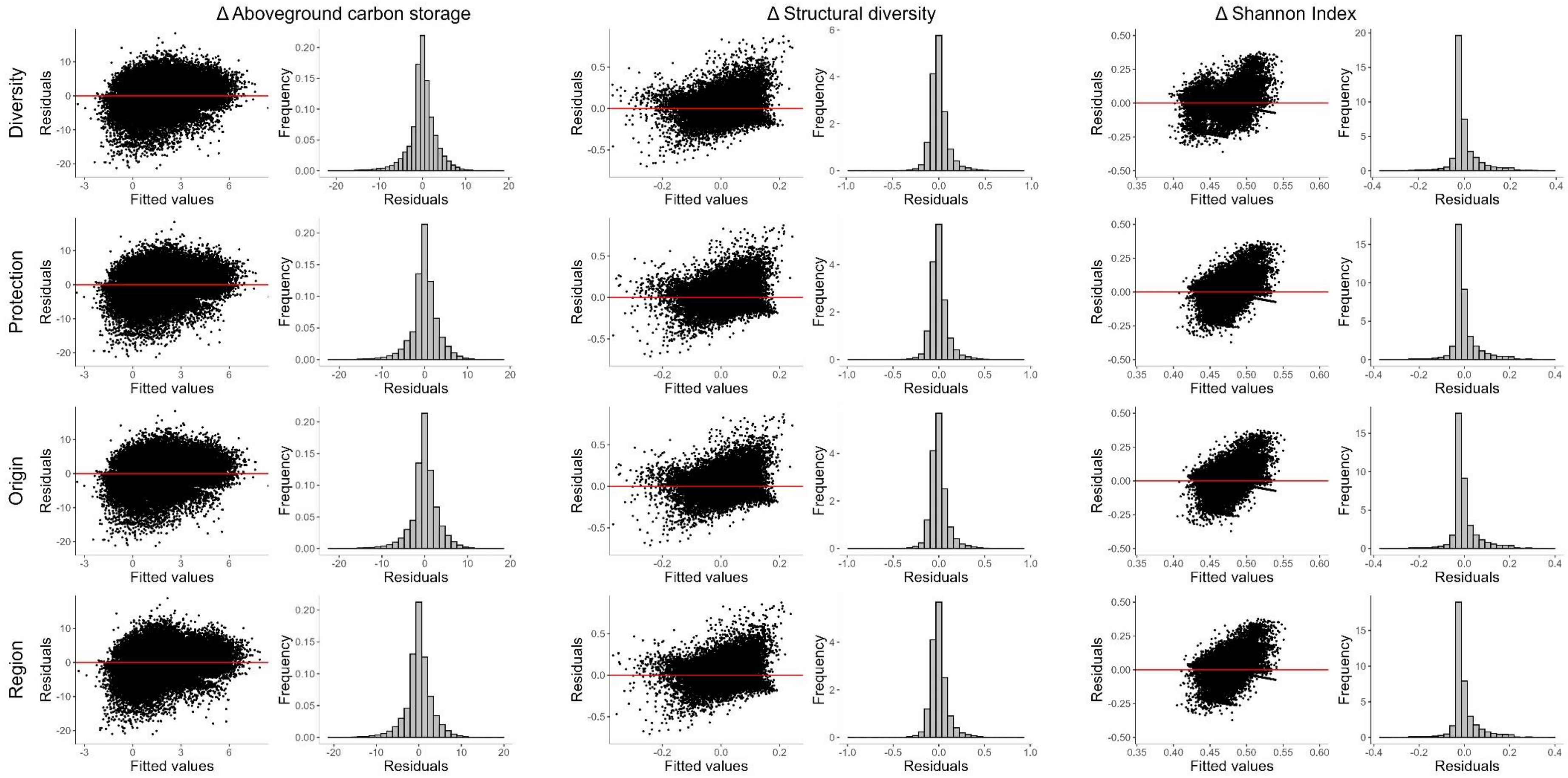

**Figure S3.2 (Cont.)**

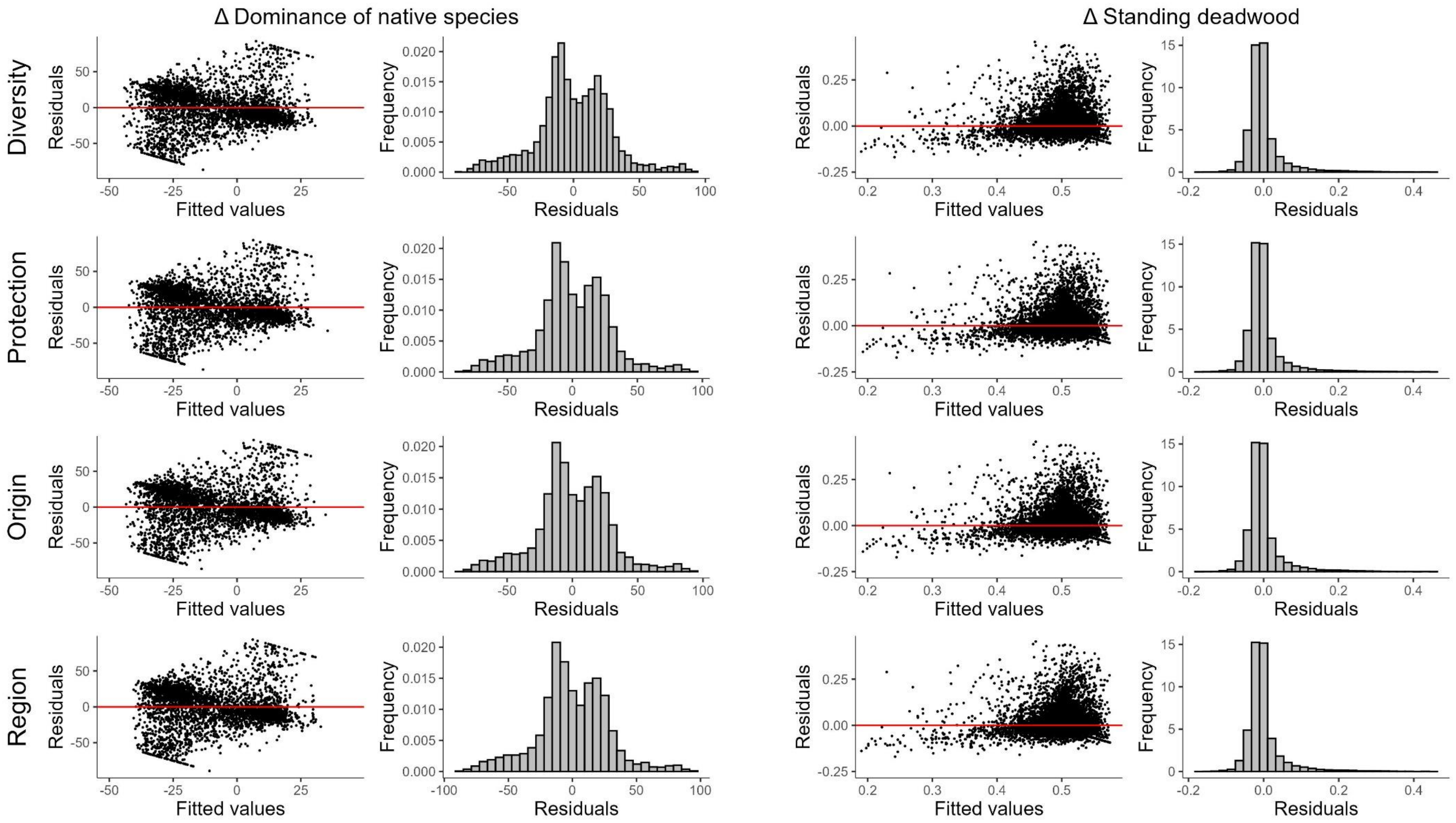

**Figure S3.3.** Scatterplot of residuals versus predicted values and histogram of the residuals resulted from mixed lineal models for changes in carbon storage, structural diversity and Shannon Index depending on the forest type and the harvesting intensity.

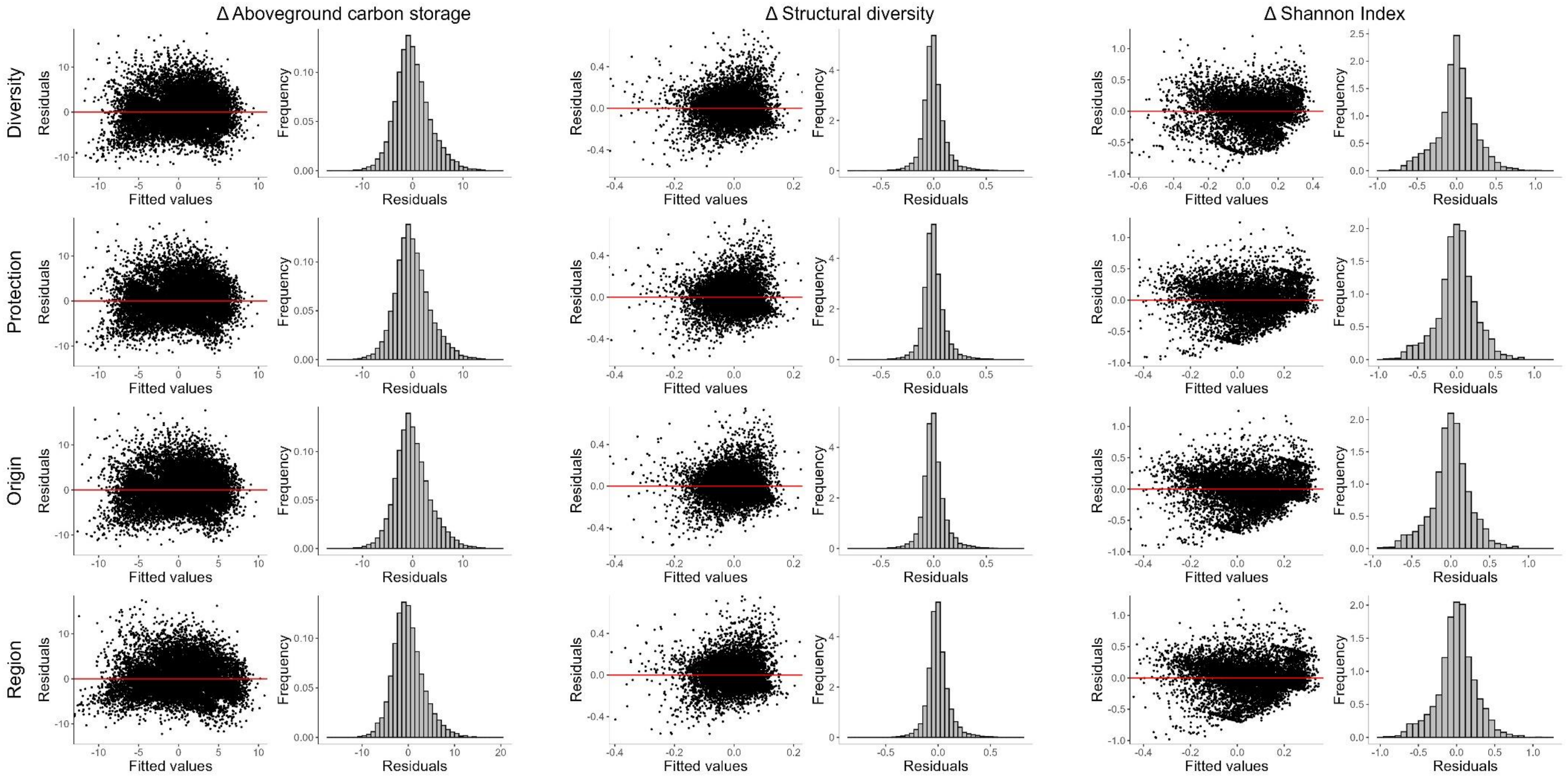

**Appendix S4. Complementary results of changes in forest conditions indicators.**

**Table S4.1**. Mean and standard deviation of forest condition indicators in each SFI census.

| | **Belowground carbon storage (Mg C ha$^{-1}$)** | | **Aboveground carbon storage (Mg C ha$^{-1}$)** | | **Structural diversity** | | **Shannon Index** | | **Dominance of native species (%)** | | **Standing deadwood (m$^2$ ha$^{-1}$)** | |
|---|---|---|---|---|---|---|---|---|---|---|---|---|
| | Mean | Std. Dev | Mean | Std. Dev | Mean | Std. Dev | Mean | Std. Dev | Mean | Std. Dev | Mean | Std. Dev |
| 2SFI | 10.55 | 18.34 | 32.29 | 41.22 | 0.30 | 0.16 | 0.18 | 0.30 | 96.90 | 16.13 | - | - |
| 3SFI | 13.47 | 20.46 | 42.81 | 50.56 | 0.31 | 0.16 | 0.21 | 0.32 | 96.29 | 17.56 | 0.37 | 1.42 |
| 4SFI | 16.47 | 23.03 | 52.53 | 59.18 | 0.33 | 0.16 | 0.25 | 0.34 | 96.33 | 17.18 | 0.68 | 1.80 |

**Table S4.2**. Mean and standard deviation of trend of forest condition indicators between censuses.

| | **Δ Belowground carbon storage (Mg C ha$^{-1}$ yr$^{-1}$)** | | **Δ Aboveground carbon storage (Mg C ha$^{-1}$ yr$^{-1}$)** | | **Δ Structural diversity yr$^{-1}$** | | **Δ Shannon Index yr$^{-1}$** | | **Δ Dominance of native species (% yr$^{-1}$)** | |
|---|---|---|---|---|---|---|---|---|---|---|
| | Mean | Std. Dev | Mean | Std. Dev | Mean | Std. Dev | Mean | Std. Dev | Mean | Std. Dev |
| 23SFI | 0.2328 | 0.6122 | 0.8913 | 2.9422 | 0.0014 | 0.0101 | 0.0034 | 0.0163 | -0.0601 | 0.7379 |
| 34SFI | 0.2338 | 0.6172 | 0.9595 | 3.3129 | 0.0013 | 0.0088 | 0.0035 | 0.0144 | -0.0203 | 0.7037 |

**Table S4.3.** Summary of the models analysing trends in biodiversity indicators for each forest classification depending on harvesting occurrence. Estimates, standard error and p values are shown for main effects and interactions.

| | **Δ Aboveground carbon storage (Mg C ha$^{-1}$)** | | | **Δ Structural diversity** | | | **Δ Shannon Index** | | |
|---|---|---|---|---|---|---|---|---|---|
| | **Diversity** | | | | | | | | |
| | Estimate | Std. Error | p | Estimate | Std. Error | p | Estimate | Std. Error | p |
| Intercept | -0.17 | 0.04 | 0.00 | -0.01 | 0.00 | 0.00 | -0.07 | 0.00 | 0.00 |
| Initial basal area | - | - | - | -0.02 | 0.00 | 0.00 | -0.02 | 0.00 | 0.00 |
| Initial tree density | 0.32 | 0.02 | 0.00 | 0.00 | 0.00 | 0.00 | -0.02 | 0.00 | 0.00 |
| Initial mean tree size | -0.29 | 0.02 | 0.00 | 0.02 | 0.00 | 0.00 | -0.01 | 0.00 | 0.00 |
| Initial indicator | -0.05 | 0.02 | 0.01 | -0.05 | 0.00 | 0.00 | -0.01 | 0.00 | 0.00 |
| Aridity index | 0.67 | 0.02 | 0.00 | 0.02 | 0.00 | 0.00 | 0.03 | 0.00 | 0.00 |
| Years between censuses | -0.01 | 0.02 | 0.36 | 0.00 | 0.00 | 0.00 | 0.01 | 0.00 | 0.00 |
| 34SFI period | 0.11 | 0.04 | 0.00 | 0.00 | 0.00 | 0.02 | 0.01 | 0.00 | 0.01 |
| Disturbances[Yes] | 0.78 | 0.03 | 0.00 | 0.01 | 0.00 | 0.00 | 0.02 | 0.00 | 0.00 |
| Mixed | -0.19 | 0.06 | 0.00 | 0.02 | 0.00 | 0.00 | -0.20 | 0.01 | 0.00 |
| Monospecific:Unharvested | 2.60 | 0.04 | 0.00 | 0.02 | 0.00 | 0.00 | -0.01 | 0.00 | 0.01 |
| Mixed:Unharvested | 2.90 | 0.07 | 0.00 | 0.02 | 0.00 | 0.00 | 0.13 | 0.00 | 0.00 |
| Conditional $R^2$ | | **0.21** | | | **0.24** | | | **0.30** | |
| Marginal $R^2$ | | **0.16** | | | **0.22** | | | **0.19** | |
| | **Protection** | | | | | | | | |
| Intercept | 0.72 | 0.15 | 0.00 | -0.01 | 0.00 | 0.03 | -0.11 | 0.01 | 0.00 |
| Initial basal area | - | - | - | -0.02 | 0.00 | 0.00 | -0.02 | 0.00 | 0.00 |
| Initial tree density | 0.32 | 0.02 | 0.00 | 0.00 | 0.00 | 0.00 | -0.02 | 0.00 | 0.00 |
| Initial mean tree size | -0.28 | 0.02 | 0.00 | 0.01 | 0.00 | 0.00 | -0.01 | 0.00 | 0.00 |
| Initial indicator | -0.05 | 0.02 | 0.01 | -0.05 | 0.00 | 0.00 | -0.04 | 0.00 | 0.00 |
| Aridity index | 0.67 | 0.02 | 0.00 | 0.02 | 0.00 | 0.00 | 0.03 | 0.00 | 0.00 |
| Years between censuses | -0.02 | 0.02 | 0.32 | 0.00 | 0.00 | 0.00 | 0.01 | 0.00 | 0.92 |
| 34SFI period | 0.12 | 0.04 | 0.00 | 0.00 | 0.00 | 0.03 | 0.01 | 0.00 | 0.00 |
| Disturbances[Yes] | 0.78 | 0.03 | 0.00 | 0.01 | 0.00 | 0.00 | 0.02 | 0.00 | 0.00 |
| Unprotected | -0.95 | 0.15 | 0.00 | 0.00 | 0.00 | 0.62 | 00.00 | 0.01 | 0.91 |
| Protected:Unharvested | 1.50 | 0.16 | 0.00 | 0.02 | 0.01 | 0.00 | 0.00 | 0.01 | 0.94 |
| Unprotected:Unharvested | 2.69 | 0.03 | 0.00 | 0.02 | 0.00 | 0.00 | 0.02 | 0.00 | 0.00 |
| Conditional $R^2$ | | **0.21** | | | **0.24** | | | **0.29** | |
| Marginal $R^2$ | | **0.17** | | | **0.22** | | | **0.16** | |

**Table S4.3 (Cont.)**

| | **Δ Aboveground carbon storage (Mg C $ha^{-1}$)** | | | **Δ Structural diversity** | | | **Δ Shannon Index** | | |
|---|---|---|---|---|---|---|---|---|---|
| | **Origin** | | | | | | | | |
| | Estimate | Std. Error | p | Estimate | Std. Error | p | Estimate | Std. Error | p |
| Intercept | -0.23 | 0.05 | 0.00 | -0.02 | 0.00 | 0.00 | -0.12 | 0.00 | 0.00 |
| Initial basal area | - | - | - | -0.02 | 0.00 | 0.00 | -0.02 | 0.00 | 0.00 |
| Initial tree density | 0.32 | 0.02 | 0.00 | 0.00 | 0.00 | 0.04 | -0.02 | 0.00 | 0.00 |
| Initial mean tree size | -0.28 | 0.02 | 0.00 | 0.01 | 0.00 | 0.00 | -0.01 | 0.00 | 0.00 |
| Initial indicator | -0.06 | 0.02 | 0.00 | -0.05 | 0.00 | 0.00 | -0.05 | 0.00 | 0.00 |
| Aridity index | 0.67 | 0.02 | 0.00 | 0.02 | 0.00 | 0.00 | 0.03 | 0.00 | 0.00 |
| Years between censuses | -0.02 | 0.02 | 0.36 | 0.00 | 0.00 | 0.00 | 0.01 | 0.00 | 0.00 |
| 34SFI period | 0.11 | 0.04 | 0.00 | 0.00 | 0.00 | 0.07 | 0.01 | 0.00 | 0.01 |
| Disturbances[Yes] | 0.78 | 0.03 | 0.00 | 0.01 | 0.00 | 0.00 | 0.02 | 0.00 | 0.00 |
| Planted | 0.02 | 0.05 | 0.70 | 0.01 | 0.00 | 0.00 | 0.02 | 0.01 | 0.00 |
| Natural:Unharvested | 2.86 | 0.06 | 0.00 | 0.02 | 0.00 | 0.00 | -0.01 | 0.01 | 0.22 |
| Planted:Unharvested | 2.60 | 0.04 | 0.00 | 0.02 | 0.00 | 0.00 | 0.03 | 0.00 | 0.00 |
| Conditional $R^2$ | | **0.21** | | | **0.23** | | | **0.29** | |
| Marginal $R^2$ | | **0.16** | | | **0.21** | | | **0.17** | |
| | **Region** | | | | | | | | |
| Intercept | -0.01 | 0.04 | 0.76 | -0.01 | 0.00 | 0.00 | -0.10 | 0.00 | 0.00 |
| Initial basal area | - | - | - | -0.02 | 0.00 | 0.00 | -0.02 | 0.00 | 0.00 |
| Initial tree density | 0.30 | 0.02 | 0.00 | 0.00 | 0.00 | 0.00 | -0.02 | 0.00 | 0.00 |
| Initial mean tree size | -0.30 | 0.02 | 0.00 | 0.02 | 0.00 | 0.00 | -0.01 | 0.00 | 0.00 |
| Initial indicator | -0.03 | 0.02 | 0.12 | -0.05 | 0.00 | 0.00 | -0.04 | 0.00 | 0.00 |
| Aridity index | 0.70 | 0.02 | 0.00 | 0.01 | 0.00 | 0.00 | 0.03 | 0.00 | 0.00 |
| Years between censuses | -0.01 | 0.02 | 0.52 | 0.00 | 0.00 | 0.00 | 0.01 | 0.00 | 0.92 |
| 34SFI period | 0.10 | 0.04 | 0.01 | 0.00 | 0.00 | 0.20 | 0.01 | 0.00 | 0.04 |
| Disturbances[Yes] | 0.76 | 0.03 | 0.00 | 0.01 | 0.00 | 0.00 | 0.02 | 0.00 | 0.00 |
| Temperate | -0.66 | 0.07 | 0.00 | 0.02 | 0.00 | 0.00 | -0.01 | 0.01 | 0.41 |
| Mediterranean:Unharvested | 2.43 | 0.04 | 0.00 | 0.02 | 0.01 | 0.00 | 0.01 | 0.00 | 0.00 |
| Temperate:Unharvested | 3.40 | 0.06 | 0.00 | 0.01 | 0.00 | 0.00 | 0.05 | 0.01 | 0.00 |
| Conditional $R^2$ | | **0.21** | | | **0.24** | | | **0.29** | |
| Marginal $R^2$ | | **0.16** | | | **0.22** | | | **0.16** | |

**Table S4.3 (Cont.)**

| | Δ Native species (%) | | | Δ Standing deadwood ($m^2\ ha^{-1}$) | | |
|---|---|---|---|---|---|---|
| | **Diversity** | | | | | |
| | Estimate | Std. Error | p | Estimate | Std. Error | p |
| Intercept | -13.38 | 1.05 | 0.00 | 0.48 | 0.00 | 0.00 |
| Initial basal area | 1.85 | 0.77 | 0.02 | 0.02 | 0.00 | 0.00 |
| Initial tree density | -1.04 | 0.79 | 0.19 | 0.00 | 0.00 | 0.00 |
| Initial mean tree size | -1.07 | 0.68 | 0.12 | 0.00 | 0.00 | 0.17 |
| Initial indicator | -16.70 | 0.49 | 0.00 | -0.03 | 0.00 | 0.00 |
| Aridity index | -2.79 | 0.50 | 0.00 | 0.00 | 0.00 | 0.00 |
| Years between censuses | 0.91 | 0.52 | 0.08 | 0.00 | 0.00 | 0.02 |
| 34SFI period | 5.39 | 1.05 | 0.00 | - | - | - |
| Disturbances[Yes] | 4.20 | 1.13 | 0.00 | 0.01 | 0.00 | 0.00 |
| Mixed | -1.04 | 1.17 | 0.38 | 0.01 | 0.00 | 0.01 |
| Monospecific:Unharvested | -4.93 | 1.36 | 0.00 | 0.02 | 0.00 | 0.00 |
| Mixed:Unharvested | 2.54 | 1.66 | 0.12 | 0.02 | 0.01 | 0.00 |
| $R^2$ | | **0.29** | | | **0.24** | |
| | **Protection** | | | | | |
| Intercept | 11.25 | 8.06 | 0.16 | 0.47 | 0.00 | 0.00 |
| Initial basal area | 1.78 | 0.77 | 0.02 | 0.02 | 0.00 | 0.00 |
| Initial tree density | -0.97 | 0.79 | 0.22 | 0.00 | 0.00 | 0.00 |
| Initial mean tree size | -1.07 | 0.68 | 0.11 | 0.00 | 0.00 | 0.25 |
| Initial indicator | -16.45 | 0.49 | 0.00 | -0.03 | 0.00 | 0.00 |
| Aridity index | -2.79 | 0.50 | 0.00 | 0.00 | 0.00 | 0.00 |
| Years between censuses | 1.01 | 0.52 | 0.05 | 0.00 | 0.00 | 0.03 |
| 34SFI period | 5.56 | 1.06 | 0.00 | - | - | - |
| Disturbances[Yes] | 4.06 | 1.13 | 0.00 | 0.01 | 0.00 | 0.00 |
| Unprotected | -25.23 | 8.05 | 0.00 | 0.02 | 0.00 | 0.00 |
| Protected:Unharvested | -20.05 | 10.50 | 0.05 | 0.03 | 0.00 | 0.00 |
| Unprotected:Unharvested | -2.07 | 1.10 | 0.06 | 0.02 | 0.00 | 0.00 |
| $R^2$ | | **0.36** | | | **0.24** | |

**Table S4.3 (Cont.)**

| | Δ Native species (%) | | | Δ Standing deadwood ($m^2$ $ha^{-1}$) | | |
|---|---|---|---|---|---|---|
| | **Origin** | | | | | |
| | Estimate | Std. Error | p | Estimate | Std. Error | p |
| Intercept | -11.73 | 1.70 | 0.00 | 0.48 | 0.00 | 0.00 |
| Initial basal area | 1.83 | 0.77 | 0.02 | 0.02 | 0.00 | 0.00 |
| Initial tree density | -0.99 | 0.79 | 0.21 | 0.01 | 0.00 | 0.00 |
| Initial mean tree size | -1.09 | 0.68 | 0.11 | 0.00 | 0.00 | 0.23 |
| Initial indicator | -16.62 | 0.49 | 0.00 | -0.03 | 0.00 | 0.00 |
| Aridity index | -2.97 | 0.50 | 0.00 | 0.00 | 0.00 | 0.00 |
| Years between censuses | 0.79 | 0.53 | 0.13 | 0.00 | 0.00 | 0.02 |
| 34SFI period | 5.66 | 1.06 | 0.00 | - | - | - |
| Disturbances[Yes] | 4.24 | 1.13 | 0.00 | 0.02 | 0.00 | 0.00 |
| Planted | -2.59 | 1.71 | 0.13 | 0.00 | 0.00 | 0.73 |
| Natural:Unharvested | 4.89 | 3.43 | 0.15 | 0.02 | 0.00 | 0.00 |
| Planted:Unharvested | -2.64 | 1.13 | 0.02 | 0.02 | 0.00 | 0.00 |
| Conditional $R^2$ | | **-** | | | **-** | |
| Marginal $R^2$ | | **0.36** | | | **0.24** | |
| | **Region** | | | | | |
| Intercept | -7.61 | 2.36 | 0.00 | 0.48 | 0.00 | 0.00 |
| Initial basal area | 1.81 | 0.77 | 0.02 | 0.02 | 0.00 | 0.00 |
| Initial tree density | -0.88 | 0.79 | 0.26 | 0.01 | 0.00 | 0.00 |
| Initial mean tree size | -0.88 | 0.68 | 0.20 | 0.00 | 0.00 | 0.20 |
| Initial indicator | -16.37 | 0.49 | 0.00 | -0.03 | 0.00 | 0.00 |
| Aridity index | -1.74 | 0.64 | 0.01 | 0.00 | 0.00 | 0.59 |
| Years between censuses | 0.50 | 0.56 | 0.37 | 0.00 | 0.00 | 0.01 |
| 34SFI period | 5.52 | 1.06 | 0.00 | - | - | - |
| Disturbances[Yes] | 4.08 | 1.13 | 0.00 | 0.02 | 0.00 | 0.00 |
| Temperate | -7.03 | 2.44 | 0.00 | 0.01 | 0.00 | 0.02 |
| Mediterranean:Unharvested | -5.51 | 2.69 | 0.06 | 0.02 | 0.00 | 0.00 |
| Temperate:Unharvested | -1.69 | 1.18 | 0.15 | 0.02 | 0.00 | 0.00 |
| $R^2$ | | **0.36** | | | **0.24** | |

**Table S4.4.** Summary of the models analysing trends in biodiversity indicators for each forest classification depending on the harvesting intensity. Estimates, standard error, statistical value and p values are shown for main effects and interactions.

| | **Δ Aboveground carbon storage (Mg C ha$^{-1}$)** | | | **Δ Structural diversity** | | | **Δ Shannon Index** | | |
|---|---|---|---|---|---|---|---|---|---|
| | **Diversity** | | | | | | | | |
| | Estimate | Std. Error | p | Estimate | Std. Error | p | Estimate | Std. Error | p |
| Intercept | 0.62 | 0.04 | 0.00 | -0.01 | 0.00 | 0.00 | 0.05 | 0.01 | 0.00 |
| Initial basal area | - | - | - | -0.02 | 0.00 | 0.00 | -0.02 | 0.00 | 0.00 |
| Initial tree density | -0.52 | 0.04 | 0.00 | 0.01 | 0.00 | 0.00 | -0.02 | 0.00 | 0.00 |
| Initial mean tree size | -1.17 | 0.04 | 0.00 | 0.02 | 0.00 | 0.00 | -0.01 | 0.00 | 0.01 |
| Initial indicator | -0.59 | 0.03 | 0.00 | -0.05 | 0.00 | 0.00 | -0.12 | 0.00 | 0.00 |
| Aridity index | 0.97 | 0.03 | 0.00 | 0.02 | 0.00 | 0.00 | 0.02 | 0.00 | 0.00 |
| Years between censuses | -0.02 | 0.04 | 0.65 | 0.00 | 0.00 | 0.00 | 0.01 | 0.00 | 0.19 |
| 34SFI period | 0.21 | 0.08 | 0.01 | -0.01 | 0.00 | 0.02 | 0.02 | 0.01 | 0.03 |
| Mixed | -0.49 | 0.07 | 0.00 | -0.03 | 0.00 | 0.00 | -0.04 | 0.01 | 0.00 |
| Monospecific: BA removed$^1$ | -430.23 | 3.95 | 0.00 | -0.04 | 0.12 | 0.78 | 3.67 | 0.31 | 0.00 |
| Monospecific: BA removed$^2$ | 13.30 | 3.94 | 0.00 | 0.69 | 0.12 | 0.00 | -1.27 | 0.32 | 0.00 |
| Mixed:BA removed$^1$ | -446.61 | 8.40 | 0.00 | -0.99 | 0.24 | 0.00 | -9.13 | 0.40 | 0.00 |
| Mixed:BA removed$^2$ | 3.33 | 8.07 | 0.68 | 0.45 | 0.24 | 0.07 | -2.62 | 0.40 | 0.00 |
| $R^2$ | **0.51** | | | **0.21** | | | **0.28** | | |
| | **Protection** | | | | | | | | |
| Intercept | 0.18 | 0.27 | 0.51 | -0.00 | 0.01 | 0.54 | 0.04 | 0.02 | 0.07 |
| Initial basal area | - | - | - | -0.02 | 0.00 | 0.00 | -0.02 | 0.00 | 0.00 |
| Initial tree density | -0.53 | 0.04 | 0.00 | 0.01 | 0.00 | 0.00 | -0.02 | 0.00 | 0.00 |
| Initial mean tree size | -1.17 | 0.04 | 0.00 | 0.02 | 0.00 | 0.00 | -0.01 | 0.00 | 0.01 |
| Initial indicator | -0.58 | 0.03 | 0.00 | -0.05 | 0.00 | 0.00 | -0.13 | 0.00 | 0.00 |
| Aridity index | 0.95 | 0.03 | 0.00 | 0.02 | 0.00 | 0.00 | 0.02 | 0.00 | 0.00 |
| Years between censuses | -0.01 | 0.04 | 0.75 | 0.00 | 0.00 | 0.00 | 0.01 | 0.00 | 0.07 |
| 34SFI period | 0.20 | 0.08 | 0.01 | -0.01 | 0.00 | 0.01 | 0.02 | 0.01 | 0.04 |
| Unprotected | 0.35 | 0.28 | 0.20 | 0.00 | 0.01 | 0.56 | 0.00 | 0.02 | 0.89 |
| Protected: BA removed$^1$ | -375.86 | 51.81 | 0.00 | -0.10 | 1.32 | 0.94 | -1.61 | 2.94 | 0.58 |
| Protected: BA removed$^2$ | 48.71 | 35.06 | 0.16 | 1.21 | 1.06 | 0.25 | 0.94 | 2.49 | 0.70 |
| Unprotected: BA removed$^1$ | -433.03 | 3.61 | 0.00 | -0.24 | 0.11 | 0.03 | -1.07 | 0.26 | 0.00 |
| Unprotected: BA removed$^2$ | 13.58 | 3.57 | 0.00 | 0.61 | 0.11 | 0.00 | -1.16 | 0.26 | 0.00 |
| $R^2$ | **0.51** | | | **0.20** | | | **0.22** | | |

**Table S4.4. (Cont.)**

| | Δ Aboveground carbon storage (Mg C ha$^{-1}$) | | | Δ Structural diversity | | | Δ Shannon Index | | |
|---|---|---|---|---|---|---|---|---|---|
| | **Origin** | | | | | | | | |
| | Estimate | Std. Error | p | Estimate | Std. Error | p | Estimate | Std. Error | p |
| Intercept | 0.47 | 0.06 | 0.00 | -0.01 | 0.00 | 0.00 | 0.02 | 0.01 | 0.00 |
| Initial basal area | - | - | - | -0.02 | 0.00 | 0.00 | -0.02 | 0.00 | 0.00 |
| Initial tree density | -0.53 | 0.04 | 0.00 | 0.01 | 0.00 | 0.00 | -0.02 | 0.00 | 0.00 |
| Initial mean tree size | -1.16 | 0.04 | 0.00 | 0.02 | 0.00 | 0.00 | -0.01 | 0.00 | 0.01 |
| Initial indicator | -0.58 | 0.04 | 0.00 | -0.05 | 0.00 | 0.00 | -0.13 | 0.00 | 0.00 |
| Aridity index | 0.94 | 0.03 | 0.00 | 0.02 | 0.00 | 0.00 | 0.02 | 0.00 | 0.00 |
| Years between censuses | -0.02 | 0.04 | 0.58 | 0.01 | 0.00 | 0.00 | 0.01 | 0.00 | 0.03 |
| 34SFI period | 0.21 | 0.08 | 0.01 | -0.01 | 0.00 | 0.00 | 0.01 | 0.01 | 0.10 |
| Planted | 0.06 | 0.06 | 0.36 | 0.01 | 0.00 | 0.00 | 0.03 | 0.01 | 0.00 |
| Natural: BA removed$^1$ | -458.71 | 6.45 | 0.00 | 0.06 | 0.21 | 0.79 | 0.17 | 0.57 | 0.76 |
| Natural: BA removed$^2$ | 21.96 | 6.48 | 0.00 | 0.44 | 0.21 | 0.04 | -0.71 | 0.56 | 0.21 |
| Planted: BA removed$^1$ | -420.22 | 4.26 | 0.00 | -0.37 | 0.13 | 0.00 | -1.43 | 0.29 | 0.00 |
| Planted: BA removed$^2$ | 10.04 | 4.19 | 0.02 | 0.69 | 0.13 | 0.00 | -1.21 | 0.29 | 0.00 |
| $R^2$ | **0.51** | | | **0.20** | | | **0.22** | | |
| | **Region** | | | | | | | | |
| Intercept | 0.42 | 0.05 | 0.00 | -0.01 | 0.00 | 0.00 | 0.04 | 0.01 | 0.00 |
| Initial basal area | - | - | - | -0.02 | 0.00 | 0.00 | -0.02 | 0.00 | 0.00 |
| Initial tree density | -0.50 | 0.04 | 0.00 | 0.01 | 0.00 | 0.00 | -0.02 | 0.00 | 0.00 |
| Initial mean tree size | -1.13 | 0.04 | 0.00 | 0.02 | 0.00 | 0.00 | -0.01 | 0.00 | 0.01 |
| Initial indicator | -0.60 | 0.03 | 0.00 | -0.05 | 0.00 | 0.00 | -0.13 | 0.00 | 0.00 |
| Aridity index | 0.84 | 0.05 | 0.00 | 0.01 | 0.00 | 0.00 | 0.01 | 0.00 | 0.02 |
| Years between censuses | 0.10 | 0.04 | 0.01 | 0.01 | 0.00 | 0.00 | 0.01 | 0.00 | 0.03 |
| 34SFI period | 0.09 | 0.08 | 0.21 | -0.01 | 0.00 | 0.00 | 0.01 | 0.01 | 0.07 |
| Temperate | 0.72 | 0.10 | 0.00 | 0.03 | 0.00 | 0.00 | 0.02 | 0.01 | 0.14 |
| Mediterranean: BA removed$^1$ | -362.75 | 4.22 | 0.00 | -0.35 | 0.14 | 0.02 | -0.58 | 0.37 | 0.12 |
| Mediterranean: BA removed$^2$ | 40.51 | 4.08 | 0.00 | 1.06 | 0.14 | 0.00 | 0.03 | 0.36 | 0.93 |
| Temperate: BA removed$^1$ | -571.29 | 6.10 | 0.00 | 0.16 | 0.18 | 0.40 | -1.14 | 0.39 | 0.00 |
| Temperate: BA removed$^2$ | -34.13 | 6.30 | 0.00 | -0.27 | 0.18 | 0.15 | -2.32 | 0.39 | 0.00 |
| $R^2$ | **0.54** | | | **0.20** | | | **0.22** | | |

**Figure S4.1.** Boxplots of forest condition indicators across the three consecutive Spanish Forest Inventory censuses (i.e. for the second, third and fourth: 2SFI, 3SFI and 4SFI). Inset letters represent groups with significant differences ($p < 0.05$) between each census. Outliers are not shown.

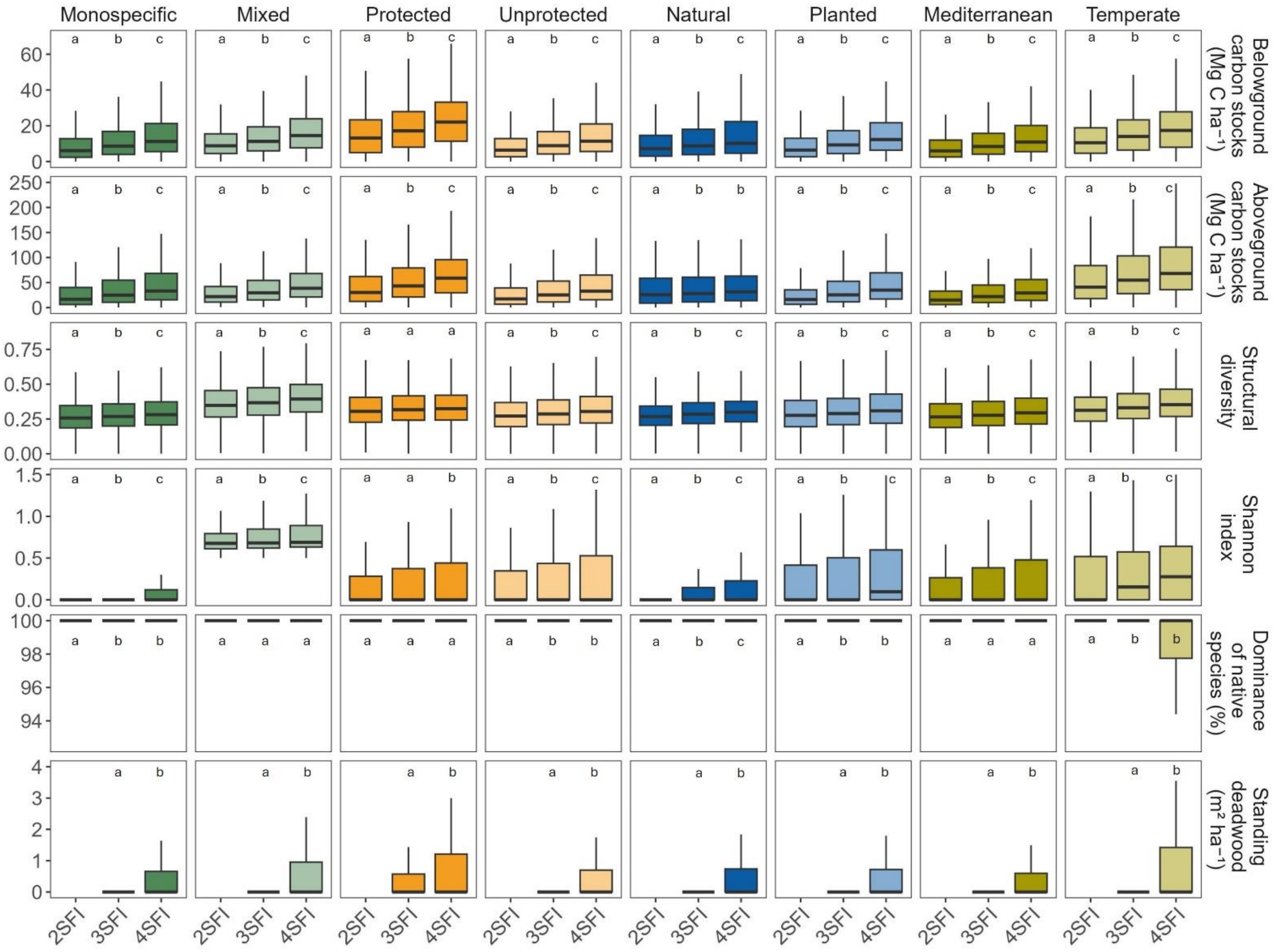

**Figure S4.2**. Boxplots of the observed trends in forest condition indicators per year between SFI periods (i.e. 23SFI and 34SFI). Inset letters represent groups with significant differences ($p < 0.05$) between each period. Outliers are not shown.

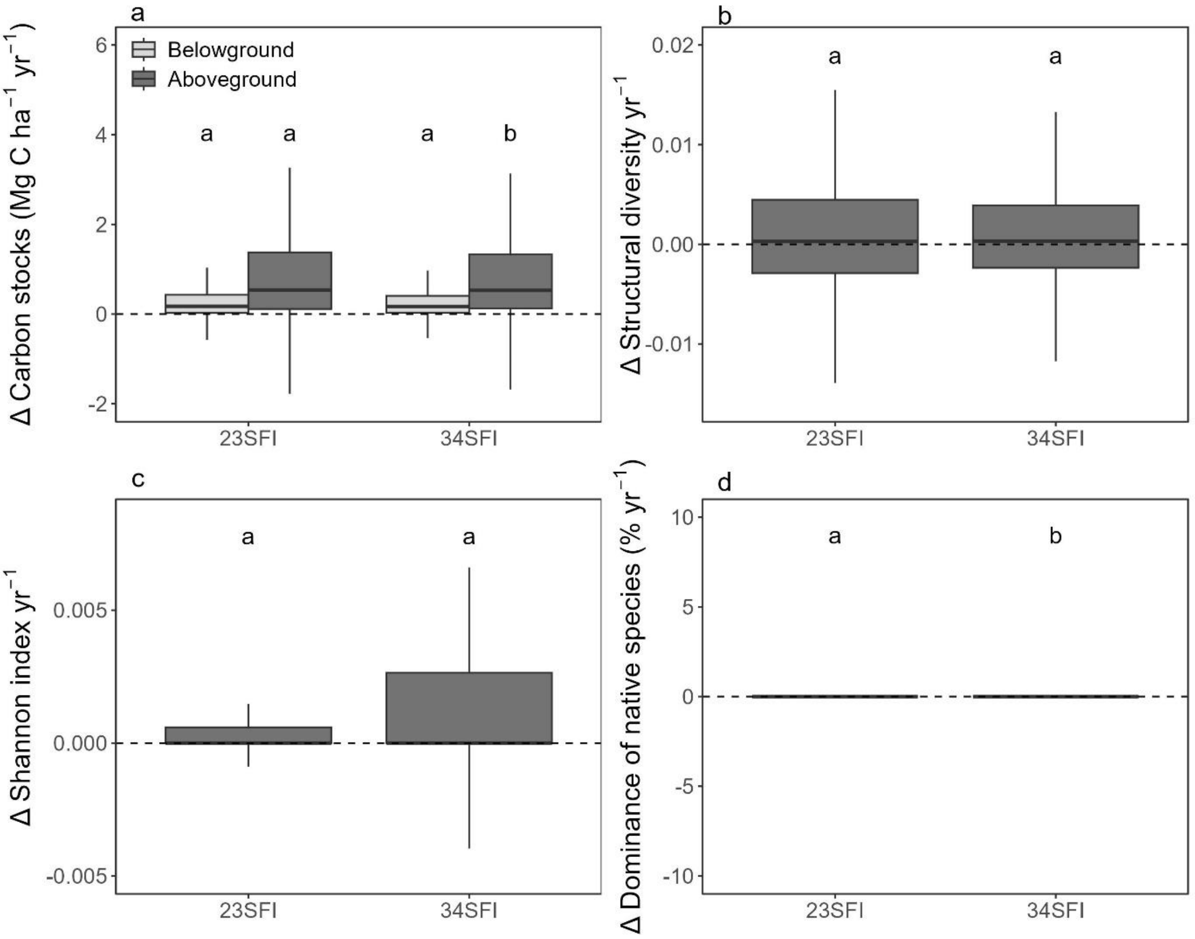

**Figure S4.3**. Boxplot of the dominance of native species (%) across the three consecutive Spanish Forest Inventory censuses (i.e. for the second, third and fourth: 2SFI, 3SFI and 4SFI). Inset letters represent groups with significant differences ($p < 0.05$) between each census.

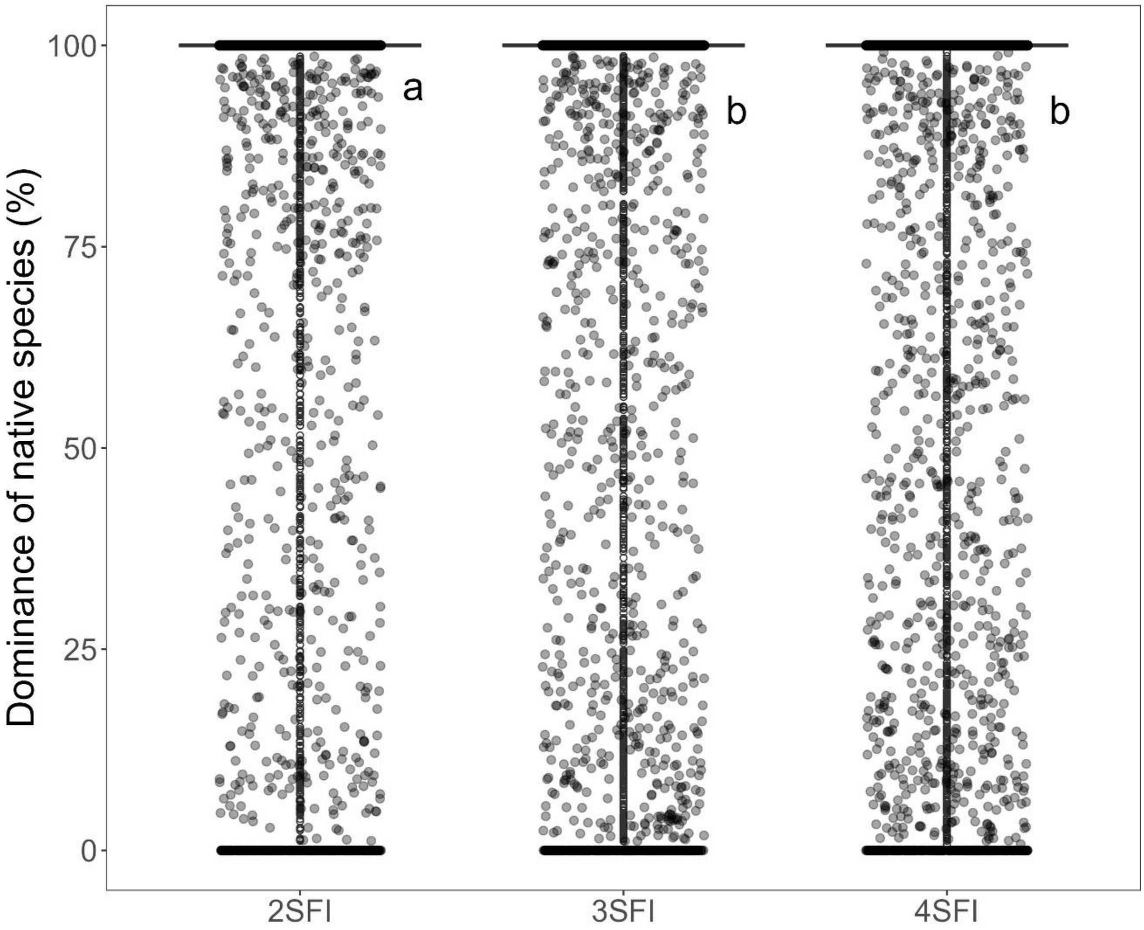
